# Simulated ischemia in live cerebral slices is mimicked by opening the Na^+^/K^+^ pump: clues to the generation of spreading depolarization

**DOI:** 10.1101/2024.09.19.613937

**Authors:** Danielle Kim, Peter Gagolewicz, Sydney McQueen, Hannah Latour, Kaitlyn Tresidder, Cathryn R. Jarvis, R. David Andrew

**Affiliations:** Dept. Biomedical & Molecular Sciences, Queen’s University, Kingston Ontario, Canada K7L 3N6

**Author notes:** Corresponding author: R. David Andrew.

**Keywords:** stroke, migraine, hemorrhage, oxygen-glucose deprivation

## Abstract

The gray matter of the higher brain undergoes spreading depolarization (SD) in response to the increased metabolic demand of ischemia, promoting acute neuronal injury and death. The mechanism linking ischemic failure of the Na^+^/K^+^ ATPase (NKA) to the subsequent onset of a large inward current driving SD in neurons has remained a mystery because blockade of conventional channels does not prevent SD nor ischemic death. The marine poison palytoxin (PLTX) specifically binds the NKA transporter at extremely low concentrations, converting it to an open cationic channel, causing sudden neuronal Na^+^ influx and K^+^ efflux. Pump failure and induction of a strong inward current should induce dramatic SD-like activity. Indeed,1-10 nM PLTX applied to live coronal brain slices induces a propagating depolarization remarkably like SD induced by oxygen/glucose deprivation (OGD) as revealed by imaging. This PLTX depolarization (PD) mimicked other effects of OGD. In neocortex, as the elevated LT front passed by an extracellular pipette, a distinct negative DC shift was recorded, indicating cell depolarization, whether induced by OGD or by bath PLTX. Either treatment induced strong SD-like responses in the same higher and lower brain regions. Further, we imaged identical real-time OGD-SD or PD effects upon live pyramidal neurons using 2-photon microscopy. Taken together, these findings support our proposal that, like most biological poisons, PLTX mimics (and takes advantage of) a biological process,ie is brain ischemia. An endogenous PLTX-like molecule may open the NKA to evoke Na^+^ influx/K^+^ efflux that drive SD and the ensuing neuronal damage in its wake.

**New and Noteworthy:** With stroke, traumatic brain injury, or sudden cardiac arrest, there is no therapeutic drug to aid brain protection and recovery. Within 2 minutes of severe ischemia, a wave of spreading depolarization (SD) propagates through gray matter. More SDs arise over hours, expanding injury. This period represents a therapeutic window to inhibit recurring SD and reduce damage but we do not understand the molecular sequence. Here we argue for a novel molecule to target.

## Introduction

Stroke is the world’s second leading cause of mortality, with an estimated lifetime risk of 8-10% (Kuriakose and Xiao 2020a). Stroke encompasses a heterogeneous group of conditions including ischemic stroke (from vessel occlusion), intracerebral and subarachnoid hemorrhage as well as cerebral venous sinus thrombosis (Bretón et al. 2012). Occlusive stroke accounts for 85% of all strokes(Kuriakose & Xiao, 2020; Woodruff et al., 2011), over half of which are attributed to large-artery ischemia ((Moustafa and Baron 2008). The remaining 15% are hemorrhagic (Bretón et al. 2012).

Focal ischemia results from a gradient of hypo-perfusion surrounding the areas supplied by the dysfunctional vessel(s). The ischemic “core” describes the region with severely low cerebral blood flow (CBF) (20% of normal levels or lower) (Hossmann 1994b) and low cerebral blood volume (CBV) with resulting low rates of oxygen and glucose turnover(Marchal et al. 1999). Here affected tissue undergoes terminal depolarization and irreversible damage in the absence of prompt treatment. Without adequate reperfusion, SD and accompanying acute neuronal injury follows within minutes (Andrew et al., 2022a,b; Kirov et al., 2020).The brain injury and neurological deficits associated with stroke are promoted by peri-infarct depolarizations (PID) which are recurrent SDs in the penumbra that expand the core (Hartings et al. 2003). Inhibition of PIDs could be an effective and valuable neuroprotective strategy, given the window of opportunity of up to 48 hours to suppress PIDs post-stroke (Hartings et al. 2003; Andrew et al.2022a)

Glutamate excitotoxicity, defined as the cell injury and death arising from prolonged exposure to accumulating extracellular glutamate and its associated cellular ionic imbalance was thought to be responsible for ischemic brain damage. However, more recent studies have found that acute cortical damage occurs independent of glutamate and the theory has been seriously questioned(Andrew et al.2022b; Obeidat et al., 2000; Renzo et al., 2010).

Spreading depolarization is a general term that includes spreading depression, PIDs as well as anoxic depolarization as a continuum (Hartings et al. 2017). Gray matter under metabolic stress *in vivo* includes various types of strokes, global ischemia and traumatic brain injury (Dreier 2011; Hartings et al. 2017). SD can also be evoked by hyperthermia, Na^+^/K^+^ pump inhibiters, hypoxia or ischemia (Hossmann 1994a, 1994b; Leao 1947; Somjen 2004).

During SD the neuronal membrane potential undergoes a rapid depolarization lasting 0.5 to 4 seconds, with K^+^, Na^+^ and Ca^2+^ ions flowing down their concentration gradients (Somjen et al. 1992; Tanaka et al. 1997)). Within several minutes of SD, pyramidal cell dendrites swell, losing their spines and forming distinct dilations or “beads” (Davies et al. 2007; Jarvis et al. 2001a; Kirov et al. 2020a), a key indicator of ischemic damage in live brain slices (Jarvis et al. 2001a) and in vivo (Murphy et al. 2008a). Understanding SD generation is important as it provides insight on possible mechanisms to inhibit PIDs and associated neuronal injury.

The hypothalamus and brainstem are more resilient to OGD in slices than the “higher” structures of neocortex, thalamus, striatum and hippocampus (Andrew 2016; Brisson et al. 2014a). Clinical support comes from the thousands of patients whose brains are globally deprived of blood flow and enter a persistent vegetative state (PVS) where their hypothalamus and brainstem are adequately functioning following global ischemia or severe TBI (Falini et al. 1998; Luigetti et al. 2012; Wytrzes et al. 1989). Also, in intact animal studies there is a general pattern of increased susceptibility to ischemic stress in higher brain compared to brainstem (Branston et al. 1984; Bures and Buresová 1981). In live brain slices, neurons in “lower” gray matter of the hypothalamus and brainstem demonstrate merely a weak SD. That is, there is a slow depolarization in response to 10 minutes of OGD, with repolarization upon return of normoxic aCSF (Brisson et al. 2012, 2013, 2014). For example, neurons in the locus ceruleus and mesencephalic neurons of the brainstem recover between 80-100% of their membrane potential, input resistance and action potential amplitude (Brisson et al. 2014b). This striking difference in the ischemic response may be tied to regionally different Na^+^/K^+^ pump isoforms. The more ischemia-resilient 1α2 and 1α3 isoforms are found in higher proportion in the hypothalamus and brainstem (Lowry et al. 2020), while the 1α1 isoform is proportionally more abundant in rostral structures(Blanco 2005; Dobretsov and Stimers 2005).

No individual neurotransmitter agonist or antagonist (including those involving glutamate receptors) blocks or delays ischemic SD onset in most brain regions (Anderson et al. 2005; Murphy et al. 2008a). In fact synapses fail before SD onset; glutamate receptor blockade does not delay SD (Andrew et al. 2022a). A non-specific Na^+^/K+ conductance has been suggested to drive SD (Czeh et al. 1993a; Somjen et al. 2009) The conclusion is that ischemic SD is extremely difficult to stop by simply blocking standard channels underlying normal neuronal function (Andrew et al., 2022a).

### The Na^+^/K^+^ Pump

The Na^+^/K^+^ ATPase (NKA) is a protein enzyme complex that spans the plasma membrane of most cells. In each cycle, the enzyme transports Na^+^ in and K^+^ out against their respective concentration gradients using the energy released from the hydrolysis of ATP. The pump cycles between an open conformational state (for either Na^+^ or K^+^) and an “occluded” state where both gates are closed, momentarily trapping the ions being transported. The occluded state ensures that a channel-like short circuit (where both gates are open which has less than a 10^-5^ probability) cannot occur (Gadsby et al. 2009a). With one complete conformational cycle required to transport 2 K^+^ in and 3 Na^+^ ions out of the cell, the pump is limited to the transporting ∼100 ions/s (Gadsby et al. 2009a) (Fig. 1, left).

**Figure 1.**
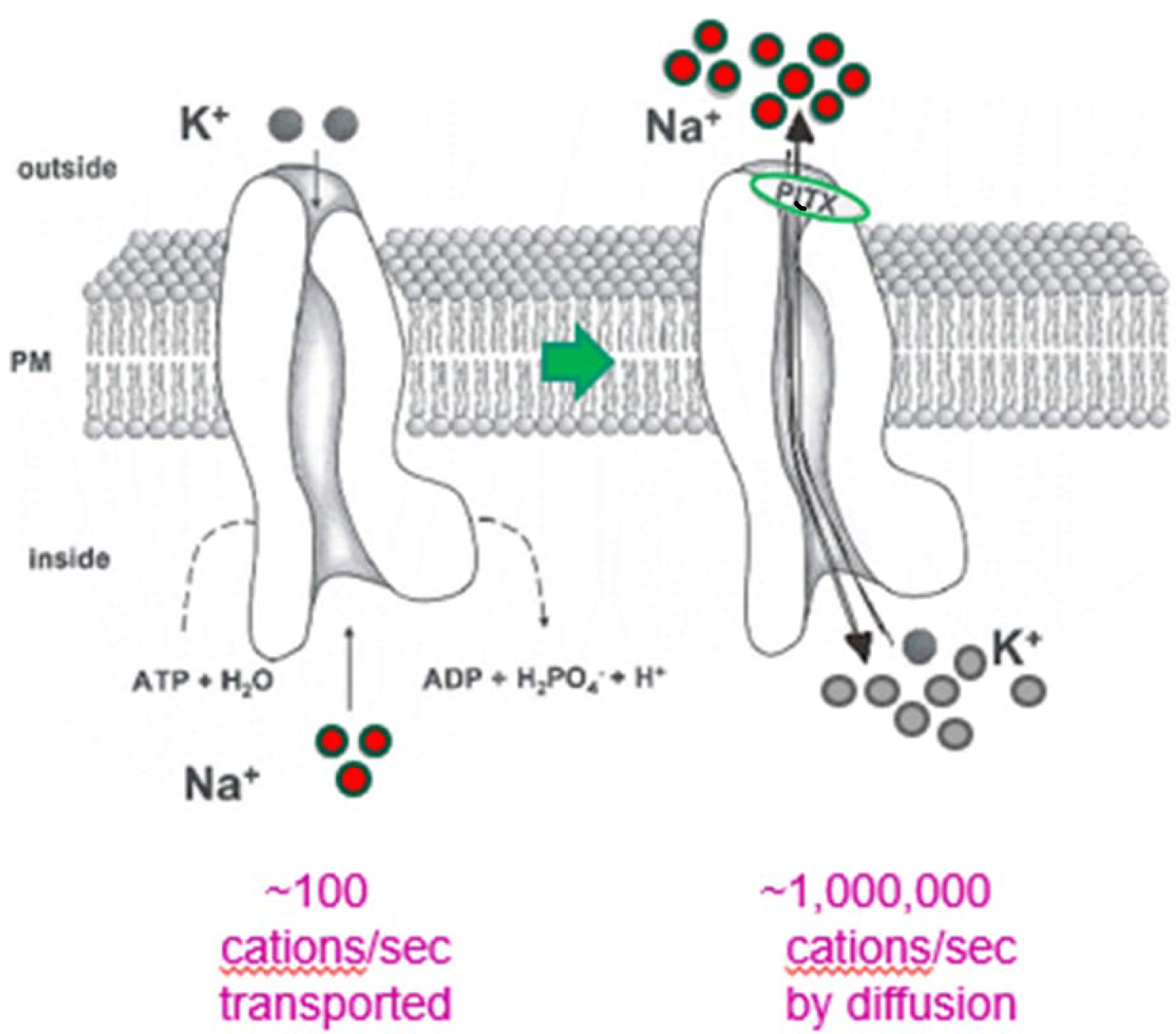
The poison palytoxin (PLTX) converts the Na^+^/K^+^ ATPase transporter (left) (Reyes and Gadsby 2006) into an open channel (right). Na^+^ and K^+^ conductances increase by orders of magnitude as they follow their concentration gradients across the plasma membrane (PM). Modified from (Gadsby et al., 2009).

The human brain expends ∼50% of available energy maintaining the Na^+^/K^+^-ATPase pump(Gottron and Lo 2009). Within 3 minutes of cardiac arrest, the ATP concentration in brain tissue decreases to 10% of normoxic values (Martin et al. 1994). Without oxygen and glucose, the pump cannot maintain the voltage and chemical gradients between neurons and extracellular space(Balestrino 1995; Martin et al. 1994). Pump failure leads to a large non-selective inward current comprising SD (Czeh et al. 1993). This current is depolarizing because Na^+^ influx exceeds K^+^ efflux (Martin et al. 1994; Tanaka et al. 1997). Importantly, it is not known how pump failure and SD are linked, nor which channel(s) conduct the formidable inward current of SD. Here we propose that SD generation involves the opening of a previously unidentified channel candidate involving the Na^+^/K^+^-ATPase (NKA).

### Palytoxin

The powerful poison palytoxin (PLTX) was originally isolated in 1961 from *palythoa toxica* corals found in Hawaii (Hilgemann 2003; Moore and Scheuer 1971). PLTX originates from dinoflagellates which are plankton(Riobó & Franco, 2011; Tosteson et al., 2003). It is one of the world’s most potent toxins(Deeds et al. 2011) detectable up the food chain in certain corals, anemones, sea urchins, crabs and fish. PLTX is a large nonpeptide with both lipophilic and hydrophilic regions. An extremely low lethal dose ranges from 0.03 to 0.45 µg/kg, (Riobó and Franco 2011b) as measured in mice, rats, rabbits, dogs, and monkeys. In early experiments, PLTX induced neuromuscular symptoms leading to follow-up studies on its ability to cause membrane depolarization(Rossini and Bigiani 2011a) which was minimally affected by the voltage-gated sodium channel blockers tetrodotoxin and saxitoxin. Subsequent investigations using erythrocytes showed that ouabain inhibited the effects induced by PLTX and vice versa, suggesting competitive inhibition for the same receptor (Habermann and Chhatwal 1982). These workers proposed that PLTX converts the Na^+^/ K^+^ transporter (Fig. 1, left) into an open pore that allows the passive diffusion of cations (Fig. 1, right) (Chhatwal et al. 1983; Gadsby et al. 2009).A series of studies, first using yeast cells which naturally lack Na^+^/K^+^ ATPase (Scheiner-Bobis et al. 1994) and later using cell-free conditions (Hirsh and Wu 1997), provided compelling evidence that PLTX specifically and reversibly induces the formation of open single channels converted from the NKA transporter (Rossini and Bigiani 2011a). PLTX acts by binding tightly to each NKA transporter externally, transforming it into a passive, nonselective cation channel (Artigas and Gadsby 2004; Chhatwal et al. 1983; Gadsby et al. 2009a), thereby allowing passage of monovalent ions and molecules smaller than 180 Da, in the order of K^+^ ≥ Na^+^ > choline >> inositol > sucrose (Chhatwal et al. 1983).

At a saturating concentration of only 100 nM, PLTX transforms thousands of pump molecules into channels in the plasma membrane of a single cell (Gadsby et al. 2009a). Despite its powerful effect at low concentration, the pump-channel does not stay permanently open while PLTX is bound. Rather, it alternates between open and closed positions throughout the bound period (Gadsby et al. 2009a; Hilgemann 2003). The duration of time the pump-channel remains open (i.e., the open probability) is affected by ATP and other ion concentrations. For instance, the open probability is five to six-fold higher in the presence of cytoplasmic ATP than without ATP (Gadsby et al. 2009a). The effect of PLTX transforming a single NKA molecule into a channel causes a 10^6^-fold gain in dissipative cation flow. Thus, in contrast to the NKA which transports about 100 ions/s, an open pump conveys perhaps a million Na^+^ and K^+^ ions per second at resting membrane potential (Fig. 1). It is estimated that only one open pump channel could overwhelm a single red blood cell to the point of death(Gadsby et al. 2009a).

### Light Transmittance (LT) Imaging of brain slices

LT imaging measures changes in tissue translucence in real time, one component of intrinsic optical signals. The swelling of neurons and/or glial cells can be detected as a local increase in LT (Andrew and MacVicar 1994; Risher et al. 2011a) because swollen cells scatter less light as membranes become more planar and so light scatter and refraction is reduced (Fig. 2). During a depolarization caused by OGD or ouabain (discussed below), the neurons and glia that depolarize then swell by mechanisms that are still unclear (Hellas and Andrew 2021), which is detected as an increase in LT. A decrease in LT may be observed within minutes in dendritic regions (Obeidat et al. 2000a) because the dendritic beads that form are the ideal diameter of 2-5 µm to scatter visible light (Malm, 1999; Andrew et al. 2007; Jarvis et al. 2001; Obeidat et al. 2000). As a real-time signal, ΔLT that can detect neuronal activation and damage, LT imaging can also be recorded simultaneously with other techniques such as patch clamping or extracellular recording. Brain slices are particularly useful when used in LT imaging, as all regions can be monitored simultaneously (Brisson et al. 2014b).

**Figure 2.**
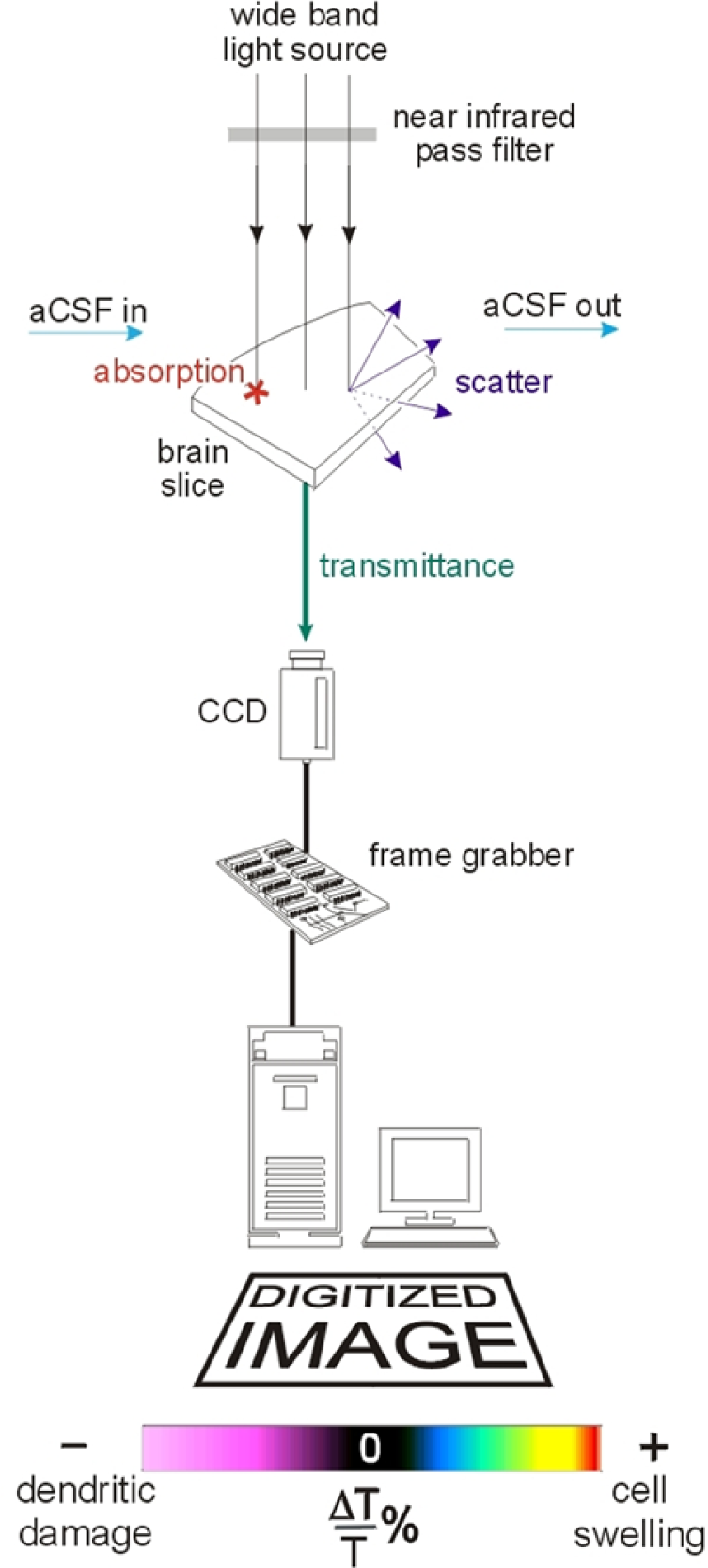
Schematic of the equipment used to image changes in light transmittance. A broadband halogen light source is directed through the brain slice. Light reaching the tissue is scattered, absorbed or transmitted. The transmitted light is collected and digitized by a charge-coupled device (CCD). Each digitized image is processed with a frame grabber board controlled by Imaging Workbench 6.0 software. Image modified from (Anderson and Andrew 2002) except that an infrared filter was not used in this study.

### The Negative Shift

One key characteristic of SD onset is the sudden negative voltage shift recorded in the extracellular space. The shift ranges from 15-35 mV *in vivo* and much less in slices and is generated by the simultaneous depolarization of many neurons near the recording electrode (Somjen et al. 1992). As the depolarizing wave front passes the recording electrode, the collective intracellular depolarization of the neurons and astrocytes generates a negative-going shift in the extracellular potential. Although the voltage may drift back to baseline, the evoked field potential is often permanently lost in the wake of SD in slices as the result of neuronal damage following OGD (Jarvis et al. 2001a).

### Two-photon laser scanning microscopy (2-PLSM)

2-PLSM enables high-resolution and high-contrast fluorescence imaging of live neurons. It affords the real-time ability to visualize swelling somata as well as dendritic spines which disappear as they swell into dendritic beads (Andrew et al., 2007; Davies et al., 2007; Risher et al., 2014)(Kirov et al. 2020). Optical sections of a specimen generate a three-dimensional reconstruction of the arborization. 2PLSM can therefore quantify the extent of damage of neurons at the subcellular level as it develops over time (Douglas et al. 2011).

### Study Objectives

The molecular mechanism linking failure of the NKA caused by ischemia to the onset of a large inward SD current in neurons is still unknown. Conventional voltage- and ligand-gated channel blockers do not prevent ischemic SD (Andrew et al., 2022a), but its rapid trajectory to near-zero millivolts points to a channel of high conductance and of high number, or of both. Furthermore, some drugs are effective in blocking or delaying SD without affecting baseline excitability (White et al. 2012), which may mean that the currents driving SD act beyond the standard channels and neurotransmitter receptors responsible for the normal physiology of neurons. The possibility of a pathological channel forming from a transporter which drives SD is plausible with the understanding of the close evolutionary relationship between transporters and channels (DeFelice & Goswami, 2007; Hirsh & Wu, 1997). Subtle changes in the pump cycle caused by lack of ATP or of inorganic phosphate (both the result of ischemia) could lead to one or both Na^+^ and K^+^ gates undergoing delayed closing or early opening (Horisberger 2004). With both gates momentarily open, the NKA essentially acts as a channel (Fig.1, right). The cation conductance results from the stabilization by PLTX of a conformation of the NKA where both gates are open, facilitated by a slowing of the closure of the external gate upon binding of K^+^, as proposed by (Horisberger 2004).

Here we ask whether SD initiation might involve the formation of a pathological channel that enables Na^+^ and K^+^ ions to rapidly flow down their concentration gradients, and if the NKA could be involved. We hypothesized that if the deadly poison PLTX causes rapid depolarization in cells by acting on the Na^+^/K^+^ pump, then exposing brain slices to PLTX could recapitulate the effects of OGD. In the present study, we set out to answer: 1) Does PLTX induce a response similar to that electrophysiologically recorded and imaged during ischemic SD? 2) Is the onset of the PLTX-induced depolarization (PD) delayed by drugs that delay ischemic SD onset? 3) Is PD weaker in lower brain gray matter vs. higher brain gray matter, as reported following simulated ischemia (Brisson et al. 2014b) or high K^+^-induced SD (Andrew et al. 2016)?

## Methods

### Brain Slice Preparation

All procedures were carried out under protocols approved by the Queen’s University Animal Care Committee. Male Sprague-Dawley (age 3-10 weeks; Charles River, St. Constant, PQ) or C57B mice of either sex (Charles River) were housed in a controlled environment (25°C, 12 h light/dark cycle) and fed Purina lab chow and water *ad libitum*. Each animal was placed in a plastic rodent restrainer (DecapiCone; Braintree Scientific, Braintree, MA) and decapitated using a guillotine. The brain was excised and immersed in ice-cold high sucrose aCSF bubbled with 95% O_2_/5% CO_2_. The composition is described below. Coronal slices (350 to 400 µm thick) of neocortex with either underlying striatum, hippocampus, or brainstem were cut using a Leica 1000-T vibratome. The slices were incubated in regular aCSF (equimolar NaCl replacing sucrose) at room temperature for 1-6 hours before being transferred to the recording chamber where they were submerged in flowing oxygenated aCSF (3 ml/min) at 32-34°C.

### Experimental Solutions

High-sucrose aCSF for dissection and slicing was composed of (in mM) 240 sucrose, 3.3 KCl, 26 NaHCO_3_, 1.3 MgSO_4_.7H_2_O, 1.23 NaH_2_PO_4_, 11 D-glucose and 1.8 CaCl_2_. Regular rat aCSF was composed of (in mM) 120 NaCl, 3.3 KCl, 26 NaHCO_3_, 1.3 MgSO_4_.7H_2_O, 1.23 NaH_2_PO_4_, 11 D-glucose and 1.8 CaCl_2_. Mouse aCSF was simply rat aCSF with the addition of 20 mM mannitol. Oxygen/glucose deprivation (OGD) aCSF which simulates ischemia *in vitro* was made by decreasing aCSF glucose from 11 to 1 mM and gassing the aCSF with 95% N_2_/5% CO_2_. A single sample of 100 ug palytoxin (Wako Chemicals USA, Richmond, VA) is supplied on a clear film in a glass vial. To make a 10 uM stock solution, 3.73 ml of ddH20 is added to the vial which dissolves the film. This yields about 3.7 ml of 10 uM palytoxin stock solution. Dibucaine or carbetapentane (Sigma-Aldrich, St. Louis, MO) were added to aCSF as required.

### Whole-Cell Patch Recording

Micropipettes were pulled from borosilicate glass (outside diameter 1.2 mm, inside diameter 0.68 mm; World Precision Instruments) to a resistance of 4-6 MΩ. The internal pipette solution contained (in mM): 130 K-gluconate, 10 KCl, 2 MgCl_2_, 1.1 EGTA, 10 HEPES, 2 Na-ATP, and 0.1 CaCl_2_ (pH adjusted to 7.3 with KOH). The junction potential was estimated to be +14 mV by using the Clampex Junction Potential Calculator. The negative resting potential measurements were not compensated for during data analysis. Visually guided whole-cell patch recordings were obtained in rat neocortical layer III pyramidal neurons. Recordings were acquired in current clamp mode of an Axoclamp-2A or 2B amplifier with a Digidata 1322 A/D converter (Axon Instruments). Clampex 10 software (Axon Instruments) was used for data acquisition with subsequent analysis using Clampfit 10 software. Sampling frequency was 10 kHz and low pass filtering was with an external Bessel filter (LPF 202a; Axon Instruments) at 2 kHz.

### Imaging Changes in Light Transmittance (ΔLT)

Imaging was used to detect the changes in LT caused by alterations in photon scattering or absorbance within live tissue in real time **(Fig. 2).** A cerebral or brainstem slice was placed in an imaging chamber with a glass coverslip base and held down with small pieces of silver wire at the slice edges. The slice was superfused with aCSF (3 ml/min) for at least 10 minutes before recording at 32-34°C. The solution superfusing the slice was switched to OGD- or PLTX-aCSF at the start of each recording session. The slice was illuminated using light from a broadband halogen light source (Carl Zeiss SNT 12V 100W) on a Zeiss upright microscope (Andrew and MacVicar 1994). The images were observed using a 2.5 X objective and the video frames were captured with a 12-bit digital camera (Hamamatsu C4742-52) at 30 Hz using Imaging Workbench 6.0 software (INDEC Biosystems, Sanata Clara, CA). The transmittance value (*T*) of the first image (*T_cont_)* was subtracted from each subsequent image (*T_expt_*) of the series, so the difference image (*T_expt_* - *T_cont_*) revealed areas where LT changed over time. To account for intrinsic regional differences in *T_cont_* (e.g. white vs grey matter), the difference signal was normalized by dividing by *T_cont_* and converted to a percentage to represent the digital intensity of the control image. Thus the change in light transmittance or ΔLT = [(*Texp - Tcont*)/ *Tcont*] x 100 = [ΔT/T] %. The ΔLT was depicted using a pseudocolor intensity scale. Changes in LT through the tissue was measured from baseline, with increased LT represented by blue-green-yellow-red pseudocoloring and decreased transmittance indicated by magenta pseudocoloring (Fig. 2). Latency to SD was the elapse time between the aCSF change and the first focal increase in LT detected prior to its propagation as SD into adjacent gray matter. Increased LT primarily indicates tissue swelling whereas decreased LT indicates dendritic beading (which scatters light reaching the detector) as noted above. As a depolarizing wave spreads through gray matter, it was imaged as a propagating front of elevated LT which is usually followed by a decrease in LT over the ensuing 10-15 minutes.

### Recording the Negative Shift

The negative DC shift is the electrical signature of sudden SD onset by cells near the recording electrode. The glass micropipette recording electrode was pulled from thin-walled capillary glass and was filled with aCSF before it was mounted on a 3-D micromanipulator. As described for LT imaging, each slice was placed in the recording chamber and superfused with aCSF at 32-34°C. The ground electrode was placed in the bath with the recording micropipette penetrating 50-100 µm deep into layer II/III of the neocortex or in CA1 stratum pyramidale of the hippocampus. Following exposure to OGD- or PLTX-aCSF, a negative DC shift in the extracellular potential representing SD was recorded as the front of elevated LT passed by the recording micropipette. The digitized data were acquired and plotted using Imaging Workbench 6.0 software (INDEC BioSystems).

### Two-Photon Laser Scanning Microscopy

For 2-PLSM, slices (350 µm thick) were prepared as above from >30 day old C57 black mice of the BG.Cg-Tg (Thy1-YFP) 16Jrs/J strain. A portion of this mouse strain’s pyramidal neurons express yellow fluorescent protein (YFP) (Feng et al. 2000), which can be visualized using 2-PLSM as detailed in (Andrew et al. 2007). The mouse slices were prepared in the same manner as rat slices but using mouse aCSF (rat aCSF + 20mM mannitol)(Joshi and Andrew 2001). The ACSF osmolarity was 292.6 mOsm and pH was 7.3–7.4. A slice was placed in the imaging chamber, held down with netting and superfused with flowing aCSF at 32-34°C. YFP^+^ neurons were imaged with a Zeiss x40 or x63 water-immersion objective lens with appropriate filter sets using a Zeiss LSM 710 NLO meta multiphoton system coupled to a Coherent Ti:sapphire laser (Brisson and Andrew 2012). Image stacks were usually taken at 3 µm increments, allowing optical sectioning of each neuronal cell body. The section with the largest area was determined visually and the cell body periphery outlined using Zeiss LSM software which was then calculated the cross-sectional area.

### Pretreatment of Slices with Dibucaine (Dib) or Carbetapentane (CP)

Slices were placed on netting in a beaker bubbled with 95% O_2_/5% CO_2_ at room temperature. Dib experiments involved pretreating slices with 10 µM or 1 µM Dib for 35-45 minutes. CP experiments involved pretreating slices with 30 µM CP for 40±5 minutes. Slices were pretreated at concentrations and pretreatment times based on previous studies(Anderson et al. 2005; Douglas et al. 2011; White et al. 2012). The onset time of OGD-SD or PLTX-induced depolarization (PD) in each drug-treated slice was calculated as a percentage of the onset time of the subsequent untreated (control) slice to accommodate for the hourly variation in SD onset time. In general, the latency to SD or PD onset slightly increased throughout each day which is typically observed in our laboratory.

### Statistical Analysis

Recordings were terminated if slices appeared visibly damaged from dissection or if slices from the same animal consistently did not display SD under OGD exposure within a window of SD onset times ranging from 4 to 8 minutes. To analyze statistical significance, unpaired t-tests and one-way repeated-measures ANOVA followed by Tukey’s post hoc method were employed using SPSS Statistics. Data were deemed statistically significant at p<0.05. All data are presented as means ± standard error.

## Results

### Whole-Cell Patch Recording

Bath application of PLTX induces depolarization in layer III pyramidal neurons (confirmed microscopically) in the neocortical slice as recorded with whole-cell patch (n=22 cells). Four different pyramidal neurons (Fig 3A) show a dose-response as superfused PLTX concentration increases from 2 to 100 nM. Responses to 50 and 100 nM reveal an SD-like response which does not recover, remaining near zero millivolts (not shown). Figure 3B shows a typical SD response to 10 minutes of oxygen-glucose deprivation (OGD) by a patched, layer III pyramidal neuron (n=25 cells). The cell did not recover its membrane potential.

**Figure 3.**
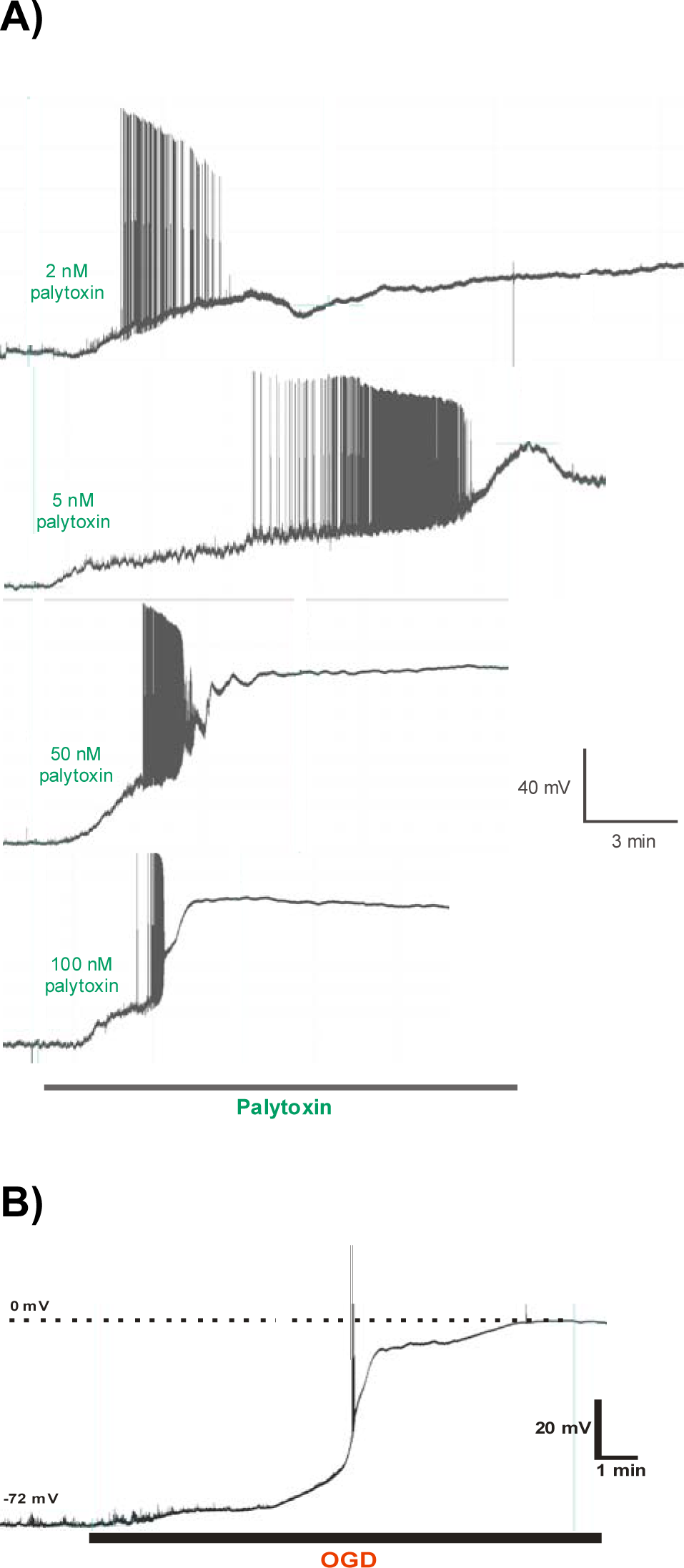
A) Palytoxin induces a spreading depolarization-like response in layer III pyramidal neurons of the neocortical slice as recorded with whole-cell patch. Four different neurons show a dose-response as superfused PLTX concentration increases from 2 to 100 nM. Responses to 50 and 100 nM do not recover. **B)** Typical SD response to 10 minutes of oxygen-glucose deprivation (OGD) by a patched pyramidal neuron.

### ΔLT Imaging of SD and of Palytoxin-induced Depolarization (PD)

ΔLT imaging revealed SD induced by 10 minutes of OGD. At SD onset, a focus of elevated LT (representing cell swelling) became a front propagating through neocortical gray matter, leaving dendritic damage in its wake which was imaged as a reduction in LT (Fig. 4). The mean (± SE) OGD-induced neocortical SD onset time was 289±10 s (n=48 slices).

**Figure 4.**
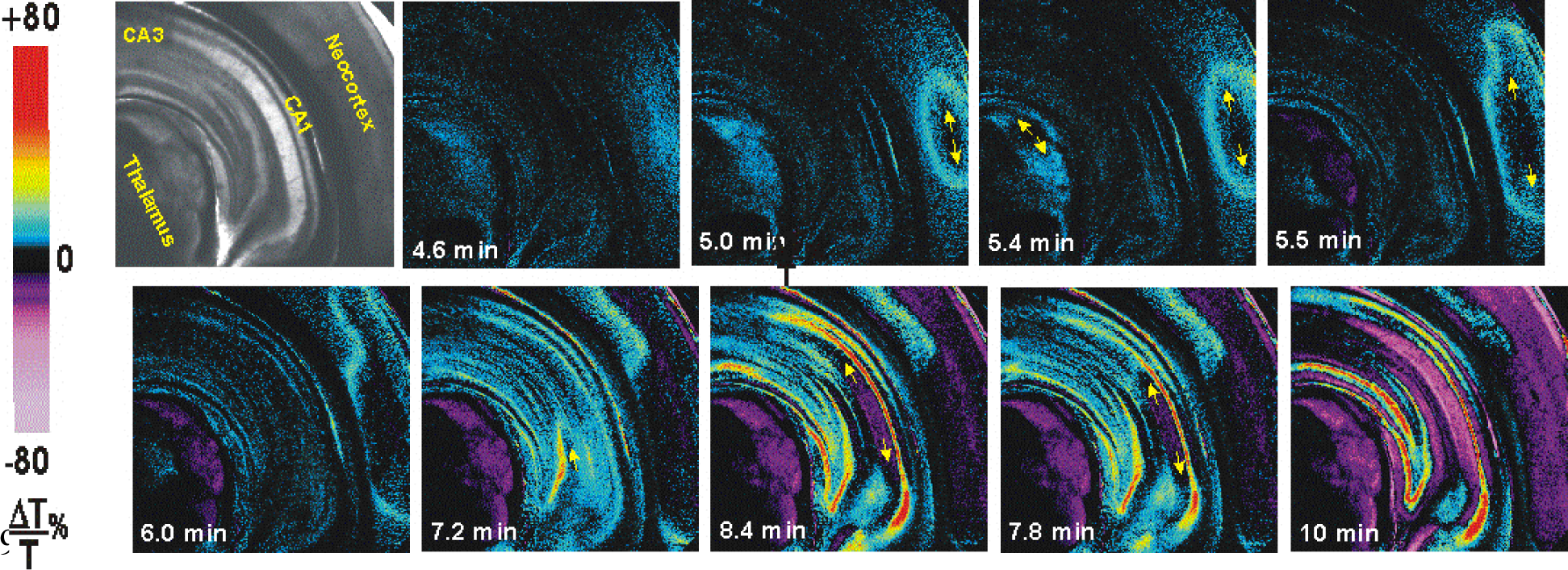
ΔLT imaging reveals damage in rat higher gray matter in the wake of SD induced by 10 minutes of OGD. The foci of elevated LT (caused by cell swelling) arises first in thalamus and then propagates as distinct waves coursing through neocortical gray, then thalamus, then CA1 of hippocampus (arrows), leaving in their wake damage that is imaged as light scatter (magenta pseudocoloring).

To help determine a low, but effective, concentration of PLTX to evoke PD in the superfused bath, 10 or 1 nM PLTX in aCSF flowed over slices for 10 minutes (slices from 8 rats). At 10 nM PLTX, the mean (± SE) neocortical PD onset time was 246 ± 14 s (n=45 slices) (Table 1). At 1 nM PLTX onset time was significantly later and more variable at 402 ± 49 s (n=16; F(2,119)=14.36, p<0.001) (Fig. 5). As with OGD, PLTX superfusion induced depolarization(s) in the hippocampus, striatum and thalamus (Table 3). The neocortex was depolarized in the highest proportion of slices in response to OGD or to PLTX. In neocortex, superfusion of 10 nM PLTX induced depolarization earlier and in a higher proportion of slices compared to 1 nM PLTX, demonstrating a dose effect (Fig. 4). Latency to depolarization in the hippocampus was significantly shorter with 10 nM PLTX compared to 1 nM PLTX (F(2,40)=13.59, p=0.04). No significant difference in the latency to depolarization between 10 nM and 1 nM PLTX was found in the striatum (F(2,41)=6.20, p=0.9) or thalamus (F(2,33)=1.42, p=0.89). To compare with PD, we also superfused OGD for 10 minutes onto slices (Fig. 4). The OGD-induced SD latency in the neocortex (289 ± 10s) was significantly shorter than the 1 nM PD latency in neocortex (p<0.001). OGD-induced SD latency was longer than the latency to 10 nM neocortical PD, but this was barely significant (p=0.09) (Table 2). The OGD-induced latencies to depolarization in the hippocampus and striatum were significantly longer than the 10 nM PD latencies induced in the same regions (p<0.001; p=0.004), while there was no significant difference between OGD and 10 nM PLTX in the latency to depolarization in the thalamus (p=0.23). Meanwhile, the latency to depolarization in the hippocampus (p=0.16), thalamus (p=0.7), or striatum (p=0.442) were not significantly different between OGD and 1 nM PLTX.

**Figure 5A,B.**
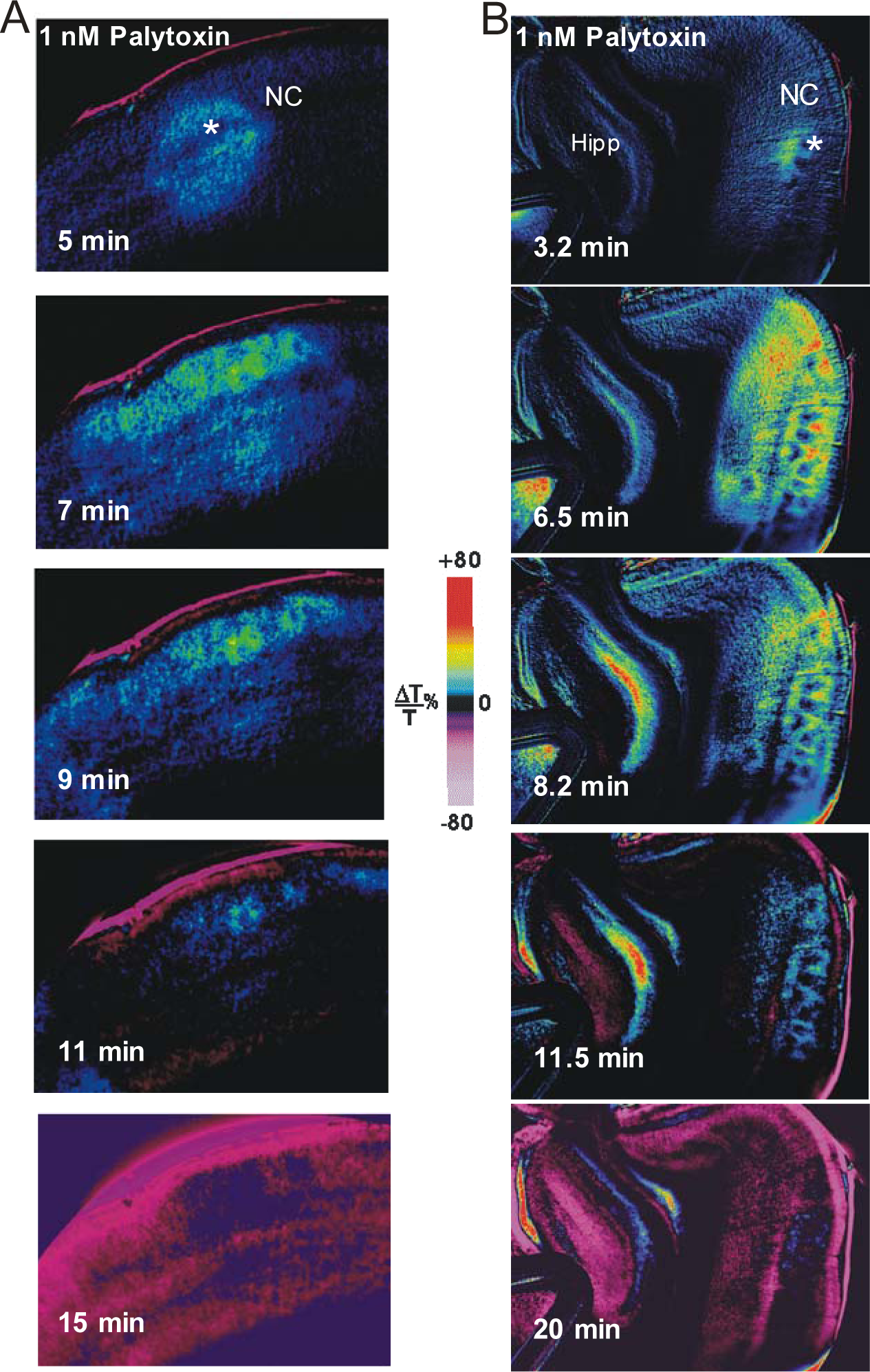
Palytoxin superfusion at 1 nM induces an OGD-like sequence of LT change in neocortex (NC). Asterisks show an initial focus of elevated LT (blue/yellow pseudocoloring) arising focally and coursing through the neocortical gray. Control aCSF is washed on after 10 minutes. The NC is subsequently damaged as shown by the reduced LT at 15 minutes (magenta pseudocoloring). Palytoxin induces OGD-like SD at extremely low concentrations by converting the Na^+^/K^+^ pump to an open cationic channel. The resulting LT front initiates and propagates (arrows) in neocortex and then hippocampus (Hipp). Note spreading swelling in the CA1 cell body layer (red) of Hipp. The regions subsequently damaged show reduced LT (magenta).

**Table 1.** Regional generation of OGD-SD or palytoxin-induced depolarization (PD) in higher brain gray matter using ΔLT imaging of superfused slices. The neocortex depolarized in the highest proportion of slices in response to OGD or to PLTX. 10 nM PLTX induced depolarization earlier (in neocortex) and in a higher proportion of slices compared to 1 nM PLTX.

|  | OGD |  |  | PLTX 10 nM |  |  | PLTX 1 nM |  |  |
| --- | --- | --- | --- | --- | --- | --- | --- | --- | --- |
|  | SD<br>(n) | SD<br>Latency<br>(s) | No<br>SD (n) | PD<br>(n) | PD<br>Latency<br>(s) | No<br>PD<br>(n) | PD<br>(n) | PD<br>Latency<br>(s) | No PD<br>(n) |
| Neocortex | 48 | 289 ± 10 | 0 | 45 | 246 ± 14 | 0 | 16 | 402 ± 49 | 2 |
| Striatum | 27 | 237 ± 16 | 2 | 15 | 158 ± 10 | 5 | 2 | 173 ± 3 | 5 |
| Hippocampus | 14 | 352 ± 27 | 6 | 20 | 175 ± 17 | 1 | 9 | 274 ± 44 | 2 |
| Thalamus | 17 | 296 ± 17 | 3 | 14 | 241 ± 24 | 5 | 5 | 263 ± 64 | 5 |

**Table 2.** Regional generation of 10 nM PLTX-induced depolarization in brainstem gray matter. The medial geniculate nucleus of the thalamus (belonging to the higher brain) and various brainstem structures displayed either a propagating signal or swelling alone in response to OGD or PLTX. The red nucleus was typical of most brainstem nuclei that showed minimal response. Superior and inferior colliculi responded only in superficial layers.

|  | OGD |  |  | PLTX |  |  |
| --- | --- | --- | --- | --- | --- | --- |
|  | n | SD | Swelling | n | SD | Swelling |
| Medial Geniculate Nuc (Thalamus) | 6 | 5 | 1 | 7 | 4 | 3 |
| Substantia Nigra Pars Reticulata | 7 | 4 | 3 | 7 | 0 | 7 |
| Superior Colliculus | 11 | 11 | 0 | 10 | 7 | 3 |
| Periaqueductal Gray | 18 | 13 |  |  | 6 |  |
| Inferior Colliculus | 8 | 8 | 0 |  |  |  |
| Red nucleus | 4 | 0 | 0 | 4 | 0 | 0 |

**Table 3.** SD or PD onset latency in slices pretreated with dibucaine or carbetapentane for 35-45 minutes prior to superfusion with OGD or PLTX. The latency to SD was compared to a subsequent untreated control slice superfused with the OGD or PLTX to determine the % delay caused by the pretreatment drug. All drug pretreatment conditions caused significant delay in SD and in PD. pretx=pretreatment.

|  | drug pretreat-<br>ment<br>(μM) | no. of<br>slices<br>(n) | drug<br>treatment<br>duration<br>(m) | treatment | mean<br>depol<br>latency<br>(s) | control vs<br>drug pretx<br>depol latency:<br>t value, p<br>value | mean %<br>delay<br>(%) | mean % delay<br>by pretx drug of<br>SD vs PD: t<br>value, p value |
| --- | --- | --- | --- | --- | --- | --- | --- | --- |
| 1 μM<br>dibucaine | control | 10 | - | OGD | 190 ± 14 | t(18)= -3.023,<br>p=0.007 | 44 ± 11 | t(17)= -0.721,<br>p=0.481 |
|  | 1 Dib | 10 | 45±5 |  | 267 ± 21 |  |  |  |
|  | control | 12 | - | PLTX | 246 ± 37 | t(22)= -1.935,<br>p=0.066 | 61 ± 20 |  |
|  | 1 Dib | 12 | 45±5 |  | 342 ± 33 |  |  |  |
| 10 μM<br>dibucaine | control | 16 | - | OGD | 200 ± 13 | t(31)= -6.303,<br>p<0.001 | 72 ± 19 | t(42)= 0.395,<br>p=0.695 |
|  | 10 Dib | 17 | 35±5 |  | 315 ± 13 |  |  |  |
|  | control | 28 | - | PLTX | 217 ± 17 | t(55)= -3.417,<br>p=0.001 | 62 ± 17 |  |
|  | 10 Dib | 29 | 35±5 |  | 286 ± 11 |  |  |  |
| 30 μM<br>CP | control | 14 | - | OGD | 206 ± 20 | t(24)= -2.801,<br>p=0.01 | 51 ± 16 | t(26)= -2.360,<br>p=0.026 |
|  | 30 CP | 12 | 40±5 |  | 288 ± 20 |  |  |  |
|  | control | 21 | - | PLTX | 207 ± 21 | t(38)= -6.402,<br>p<0.001 | 129 ± 29 |  |
|  | 30 CP | 19 | 40±5 |  | 398 ± 21 |  |  |  |

The LT imaging characteristics of PD in the neocortex and subcortical regions (Fig. 5) were strikingly similar to SD induced by OGD (Fig. 4). Both OGD- and PD-generated depolarization in the neocortex and often in the subcortical “higher” brain structures (thalamus, striatum, hippocampus). A decrease in LT was then often detected in these regions where an LT front had passed through. The decrease developed slowly over the ensuing 10-15 minutes. The maximum ΔLT was similar between OGD (n=8) and 10 nM PTX (n=8) (t(8)= -0.23, p=0.83), (Fig. 7), although PLTX displayed a greater minimum ΔLT than OGD (t(14)=2.47, p=0.03). Like OGD-SD (Fig. 4), when PD was observed in the striatum, hippocampus or thalamus, these areas also displayed decreased LT following the front of PD (Fig. 5B).

**Figure 6.**
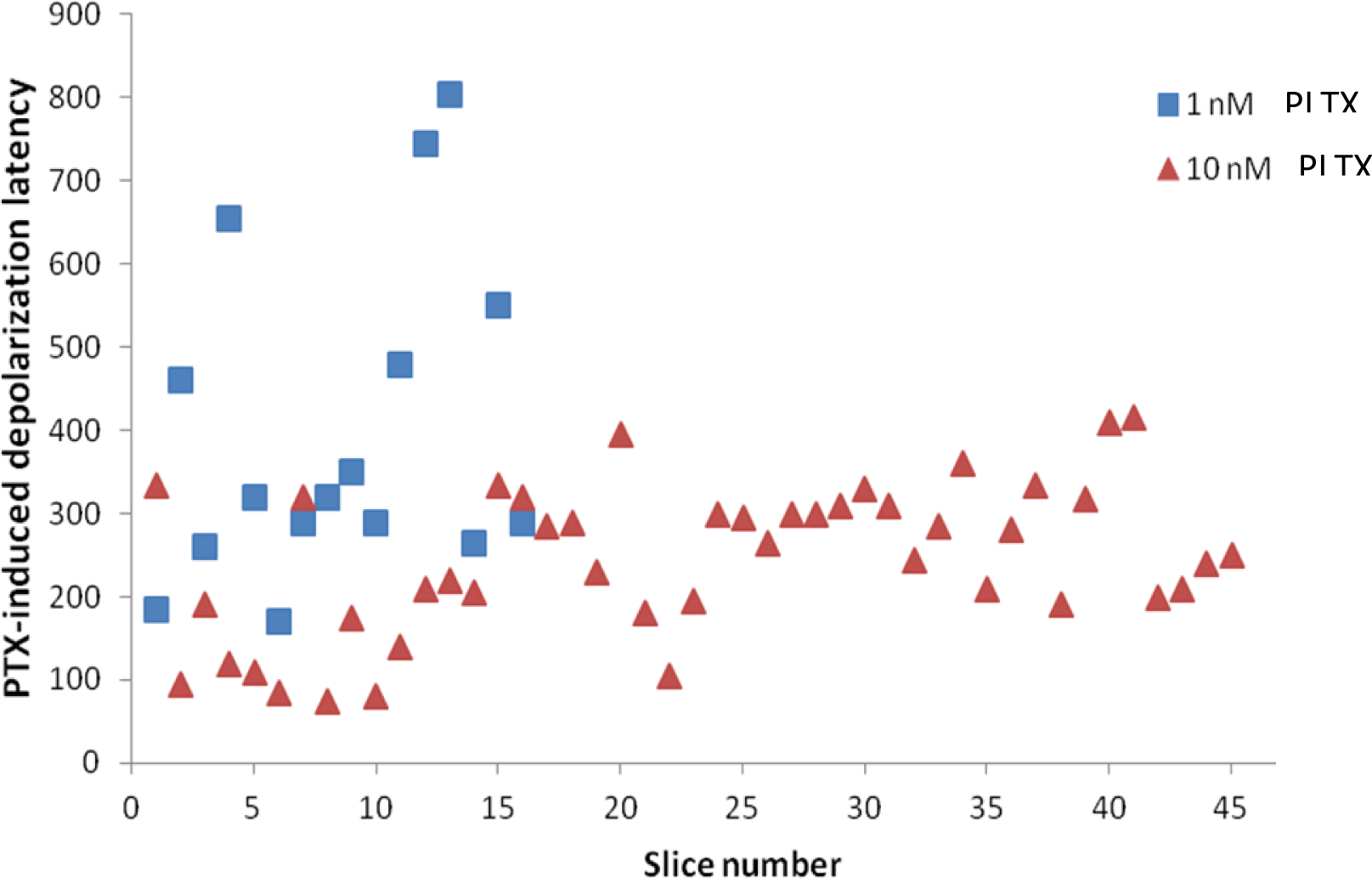
Distribution of palytoxin-induced depolarization (PD) onset latency in seconds measured by ΔLT imaging induced by superfusion of 10 or 1 nM PLTX in neocortex. At 1 nM PLTX, PD initiated later and over a wider range of time. At 10 nM PLTX, PD onset was significantly earlier (p<0.001) within a narrower time range, demonstrating a dosage effect.

**Figure 7.**
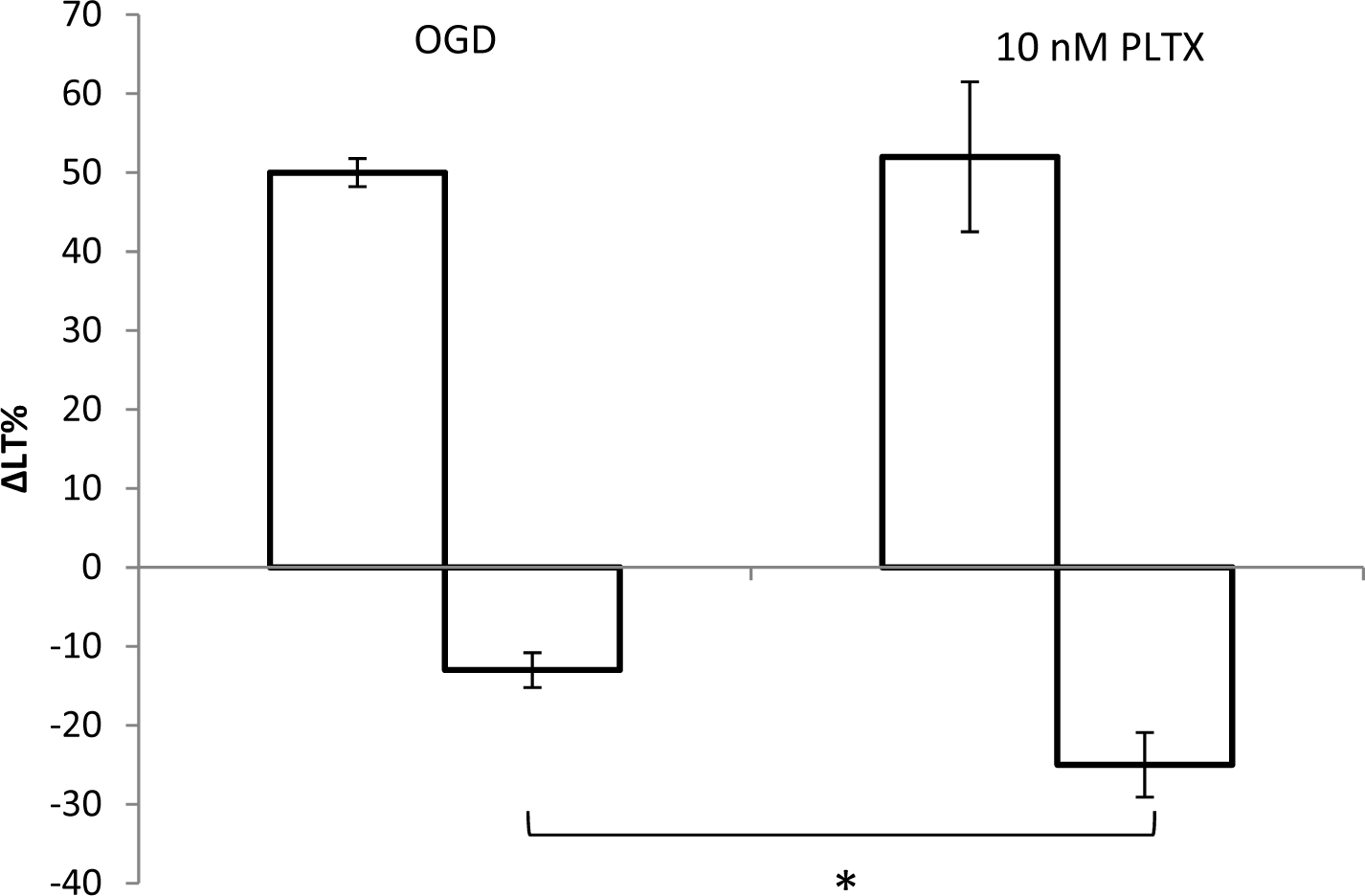
Maximum and minimum LT changes in the neocortex induced by OGD or by 10 nM PLTX in terms of spreading depolarization strength (peak LT) and subsequent injury (lowest LT value). Peak LT values were comparable between OGD and PLTX (t(8)= -0.228, p=0.826), while the LT minimum evoked by PLTX showed a significantly greater LT decrease compared to OGD (t(14)=2.474, p=0.027).

### ΔLT Imaging of OGD- and PLTX-Induced Depolarization in the Brainstem

Our imaging of SD induced by 10 minutes of OGD in the brainstem extends earlier studies in our lab examining the acute effects of OGD in brainstem (Brisson et al. 2012, 2013, 2014b). Swelling, often followed by SD, was observed in the superficial regions of superior colliculus and inferior colliculus, in periaqueductal gray and in substantia nigra reticulata (Table 2). The medial geniculate nucleus (the most caudal part of the thalamus) also usually displayed SD as expected. Similarly, brainstem slices superfused with 10 nM PLTX for 10 minutes exhibited swelling often accompanied by PD in the same brainstem regions affected by OGD (Table 3). As expected, 10 nM PLTX induced PD earlier and within a narrower range than 1 nM PLTX (Fig. 6).

The red nucleus in midbrain did not show increased LT under OGD or PLTX. Also, the substantia nigra compacta displayed no notable increase in LT before exhibiting decreased light transmittance in response to OGD or PLTX exposure (Table 2). In contrast, the substantia nigra reticulata displayed an increase, but no decrease in LT, when imaged for 10-40 minutes after exposure to 10 minutes of OGD or PLTX. These observations suggested minimal damage to the red nucleus and substantia nigra reticulata as we previously reported.

### Extracellular Recording of OGD and PLTX-Induced Depolarization

To confirm that the moving front of elevated LT represents a regional depolarization of neurons, during OGD we measured the change in the extracellular voltage within the CA1 hippocampal region while simultaneously imaging LT changes in the same region. We first obtained a baseline recording before switching to OGD-aCSF. During the first few minutes of OGD there was no change in the baseline potential. But at the moment the SD front reached the recording electrode tip (Fig. 8A, left), there was a distinct negative-going shift in the field potential (Fig. 8A, right). This sudden drop in the voltage displayed a mean amplitude of -4.2 ± 1.0 mV (n=9) for OGD. With superfusion of PLTX, the results using this simultaneous recording technique were similar (Fig. 8B). The mean voltage amplitude was -4.1±0.8 mV (n=14) during PLTX exposure (Fig. 9) which was not significantly different from OGD (t(21)=0.06, p=0.9). The key finding for OGD or PLTX exposure was the coincidence of the negative shift onset as the LT front reached the micropipette tip. Both parameters are well established indicators of SD.

**Figure 8.**
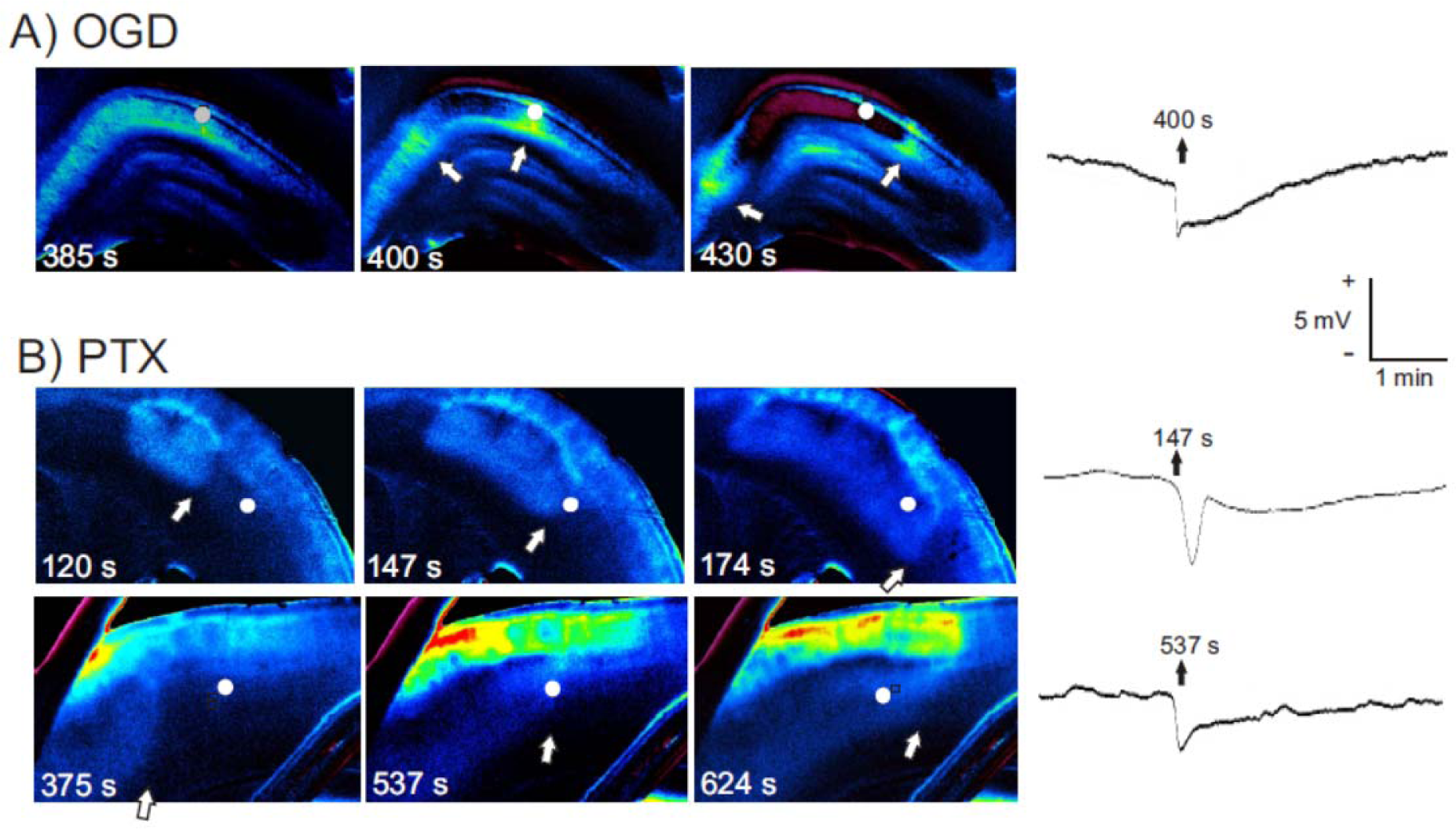
When a front of elevated LT (arrow) passes the recording electrode tip (dot in images, left), a negative DC shift is recorded (right). **A)** OGD evokes SD that propagates across the CA1 region of the hippocampus, recorded both as an elevated LT front (arrow) and a negative voltage shift as the front passes the tip of the recording micropipette (white dot). **B)** Likewise, bath application of 10 nM PLTX evokes a similar negative shift in the extracellular voltage recorded in layers II/III of the neocortex at the moment that the LT front (arrow) passes the recording pipette (white dot).

**Figure 9.**
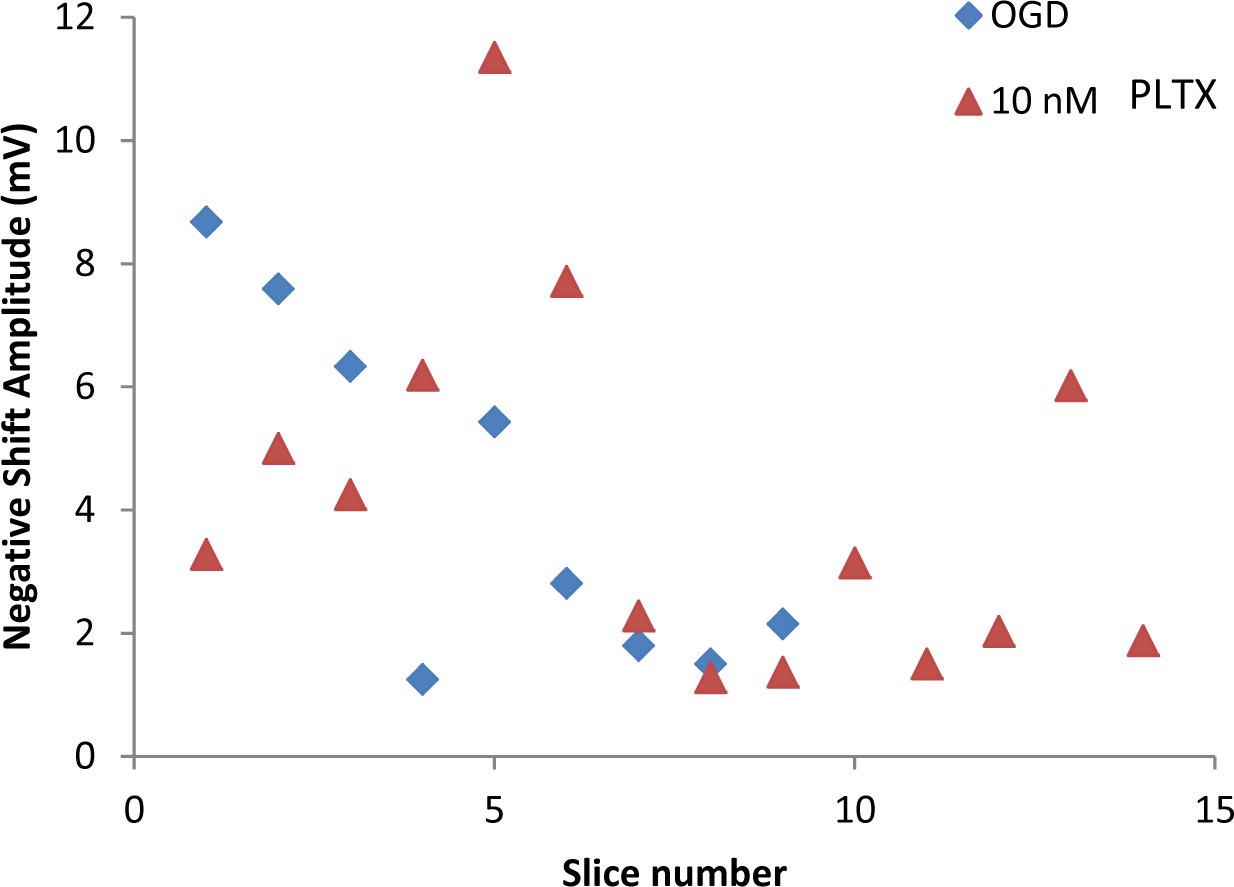
Amplitude of the negative-shift extracellular voltage during depolarization induced by OGD or by palytoxin superfusion. There is no significant difference in response between the two treatments (t(21)=0.949, p=0.709).

### Two-Photon Laser Scanning Microscopy (2-PLSM)

To confirm that, like OGD, PLTX caused cellular swelling and dendritic beading as implicated by our LT imaging, we used 2-PLSM to image live pyramidal neurons. Slices were imaged before and immediately after exposure to 10 minutes of OGD (Fig. 10) or of 10 nM PLTX (Fig. 11). In the neocortex, OGD caused a mean pyramidal cell body swelling by 53 ± 3% (n=68), calculated by measuring the largest cross-sectional area of each soma (Fig. 12). Similarly, PLTX exposure swelled pyramidal cell bodies by a mean of 47±3% (n=106) (Fig. 12). The somata swelling caused by OGD or PLTX were not significantly different (t(172)= -1.45, p=0.1). In addition, either 10 mM PLTX or OGD for 10 minutes caused a similar loss of dendritic spines as swelling progressed. The dendrites often formed chains of dilations or “beads” evoked by OGD or by PLTX.

**Figure 10A,B.**
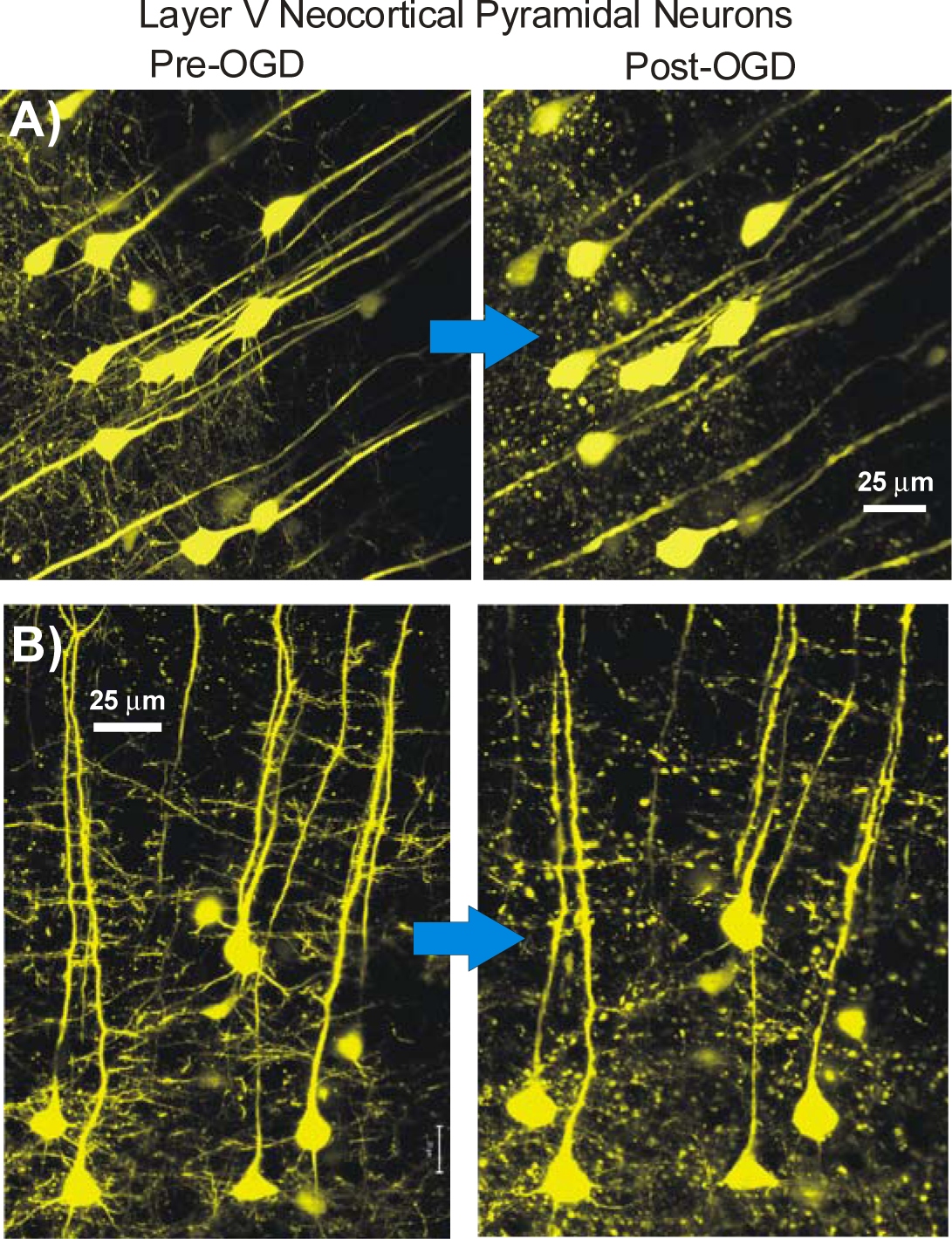
Two-photon laser scanning microscopy (2-PLSM) images of live pyramidal neurons before and after OGD. A proportion of pyramidal neurons in all regions of neocortex are YFP-positive in YFP^+^ transgenic mice. Immediately following SD induced by 10 minutes of OGD (arrows), layer V neocortical pyramidal neuron cell bodies and dendrites swell with the dendrites losing spines and becoming beaded, forming a background of fluorescent dots.

**Figure 11.**
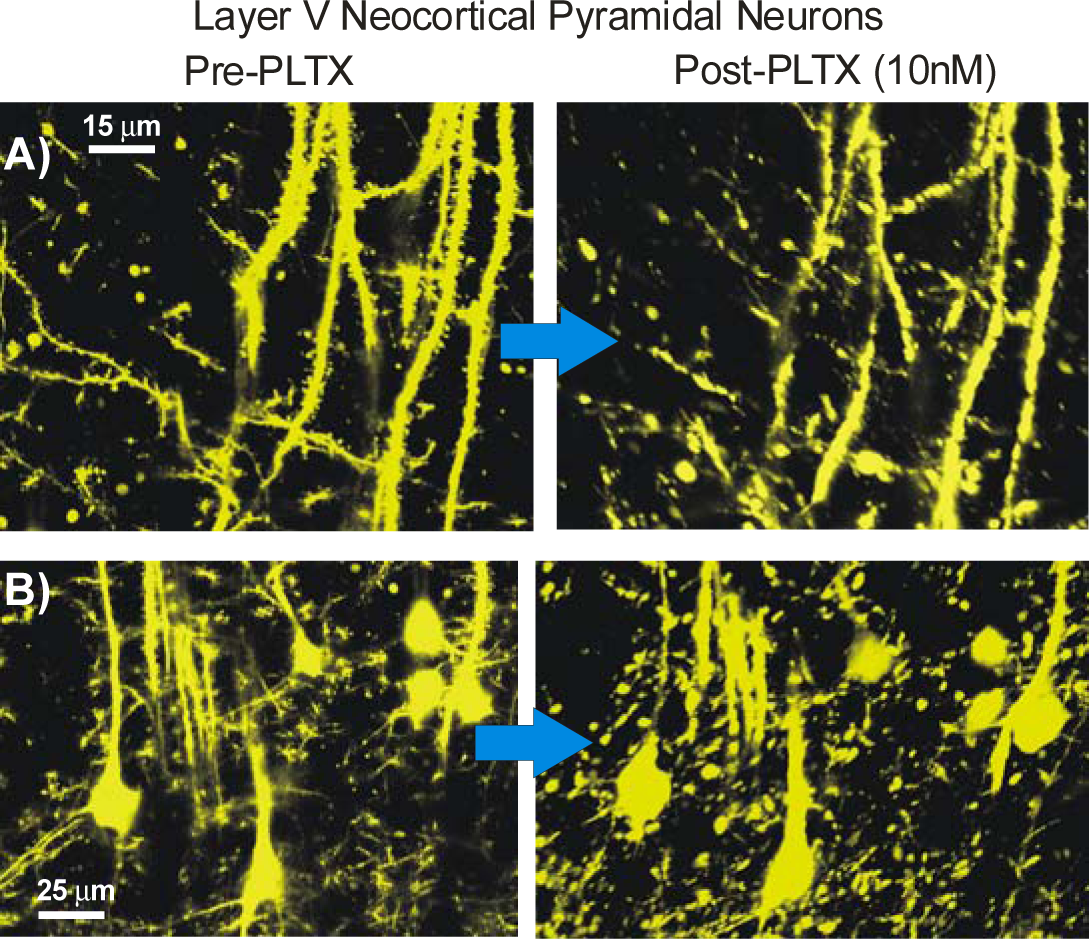
2-PLSM images of pyramidal neurons before and after 10 nM palytoxin aCSF exposure. Similar to the previous figure showing pyramidal cell responses to OGD, neocortical layer V pyramidal neuron cell bodies swell and their dendrites bead upon depolarization, induced by 10 min of PLTX. As previously detailed using OGD (Andrew et al. 2007; Risher et al. 2010), dendrites likewise swell in response to PLTX, losing their dendritic spines and forming dysmorphic and beaded dendrites.

**Figure 12.**
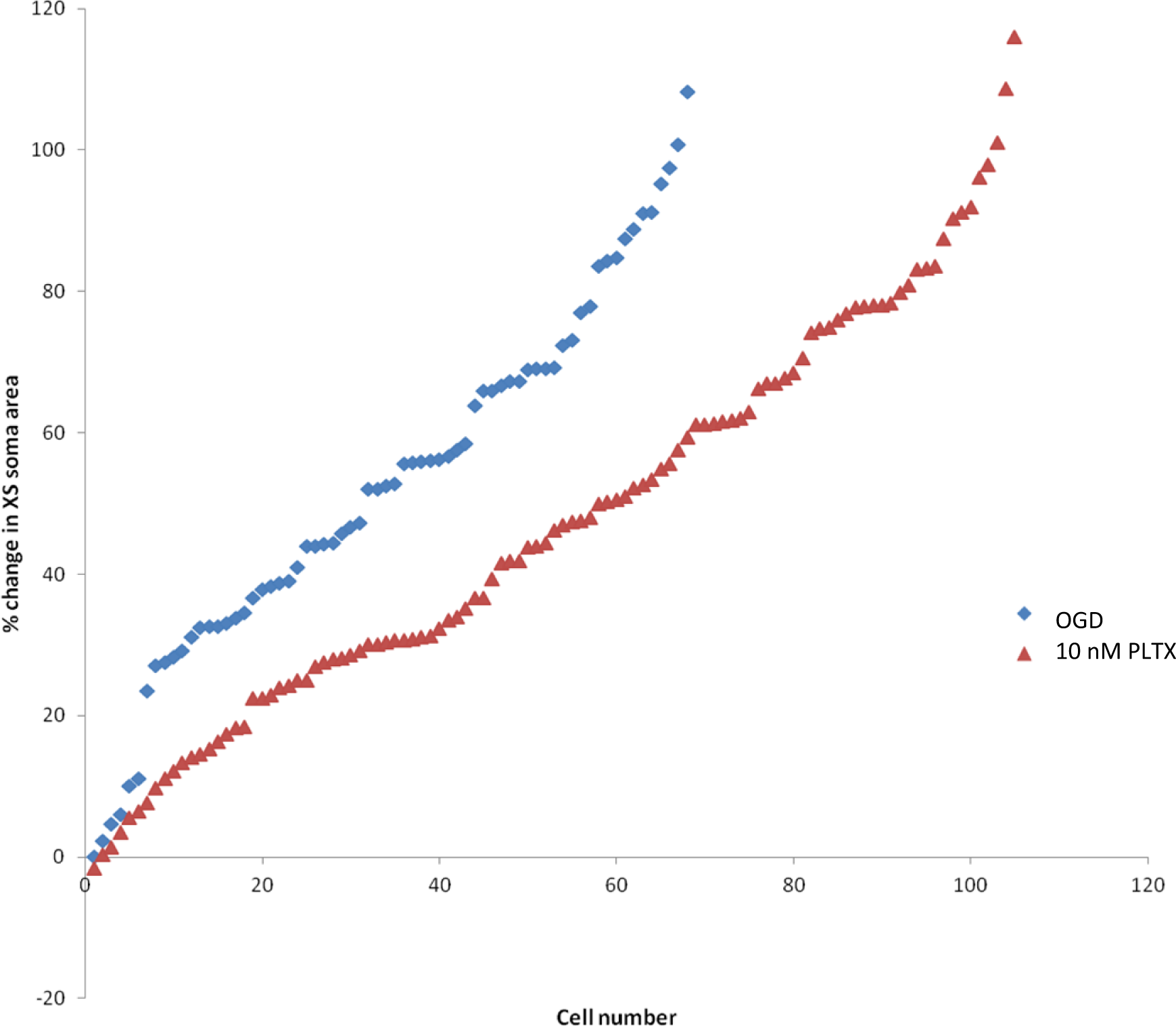
Distribution of pyramidal neuron soma swelling induced by 10 min of OGD (n=68) or by 10 nM PLTX (n=103). The change in the cross-sectional (XS) area for each cell body is plotted for both populations. Both treatments show a similar wide range (up to 110 to 115%) of irreversible swelling by pyramidal cell bodies.

### ΔLT Imaging of AD and PD in Slices Pretreated with Dibucaine and Carbetapentane

To further examine similarities between OGD and PLTX exposure, we tested two drugs previously shown to delay OGD-SD onset (White et al. 2012) to observe if either of these drugs also delay PD. Pretreatment of slices with dibucaine (DIB,, 1 or 10 μM) or carbetapentane (CP, 30 μM) caused significantly longer SD and PD onset latency compared to no pretreatment (Table 3). Pretreatment with 10 μM DIB caused a longer delay of SD than 1 μM DIB (Fig. 13). The delay in PD by 10 μM and 1 μM DIB was similar. At 30 μM, CP was more effective in delaying PD than OGD-SD (t(26)= -2.36, p=0.026). Otherwise, both drugs were similar in the delay that they imposed on either OGD-SD onset or on PD onset (Fig.12).

**Figure 13.**
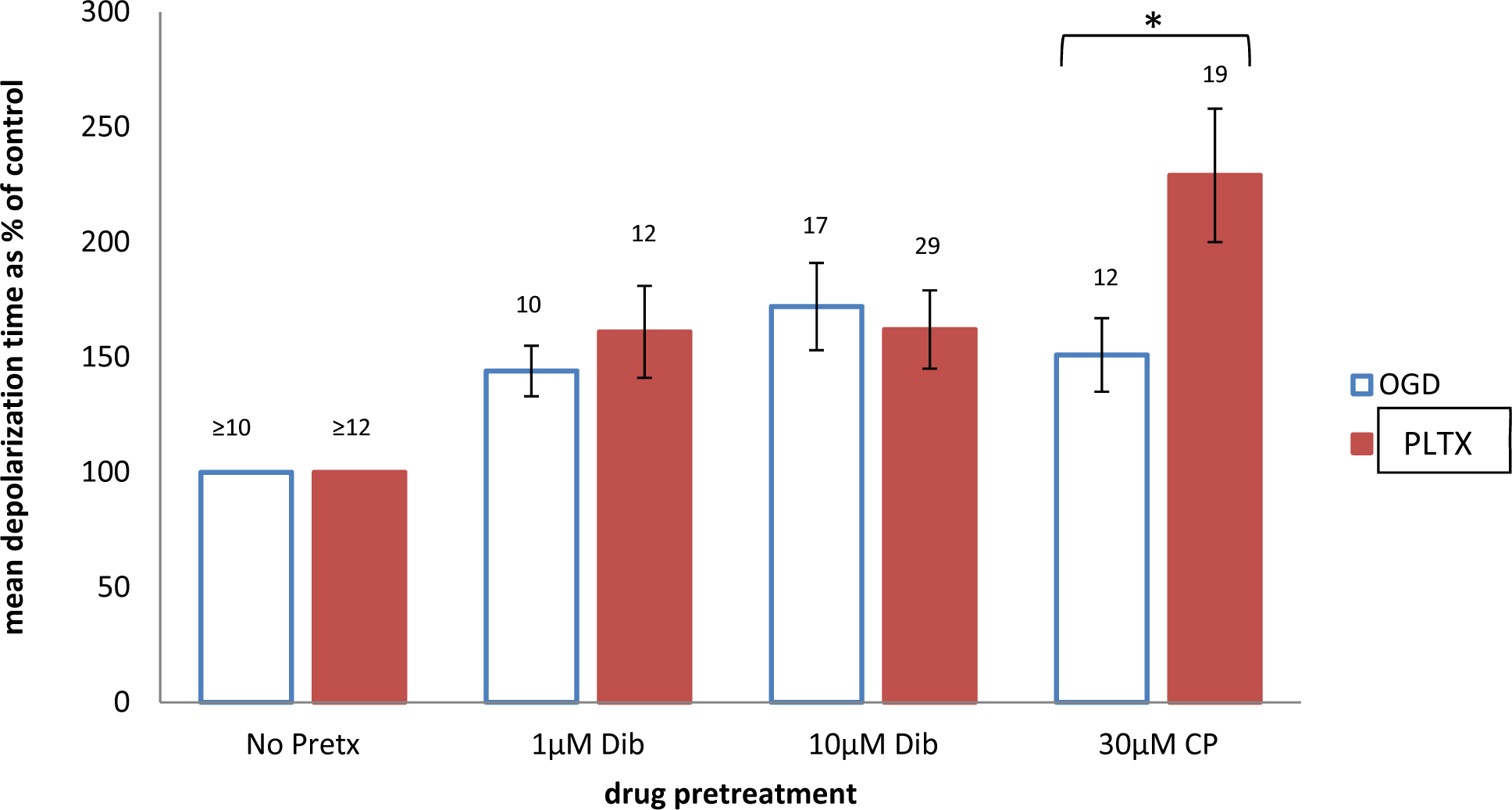
SD and PD onset latencies are similarly delayed in slices pretreated with dibucaine or carbetapentane for 35-45 minutes before superfusion with OGD aCSF or with 10 mM PLTX. The latency to depolarization of a drug-pretreated slice was compared to a subsequent untreated control slice superfused with the same PLTX or OGD to determine the % delay. 30 µM CP caused a PD delay that was significantly greater than the delay of SD, as indicated by the asterisk (*) (t(26)= -2.36, p=0.026). Otherwise, the delay between SD and PD was not significantly different following 1 µM dibucaine (t(17)= -0.72, p=0.48) or 10 µM dibucaine (t(42)=0.39, p=0.69) pretreatment.

## Discussion

Spreading depolarization is a major cause of early brain injury following ischemia (Andrew et al., 2022). A non-specific Na^+^/K^+^ conductance has been proposed to drive SD (Czeh et al. 1993) as further detailed by Somjen et al. (2009). The SD reversal potential near 0 mV supports this scenario, although the source of this TTX-insensitive inward current is still unknown. Likewise, PLTX induces a TTX-insensitive, non-specific Na^+^/K^+^ conductance by opening the NKA transporter (Artigas and Gadsby 2002, 2004, 2006; Gadsby et al. 2009a; Hilgemann 2003; Hirsh and Wu 1997; Horisberger 2004; Rossini and Bigiani 2011a).

### PTLX-induced Depolarization: Similarities with OGD-induced SD

The current study compares the effects of the potent marine poison PLTX with simulated ischemia upon mammalian gray matter. Specifically, we investigated if PLTX induces a mass propagating depolarization of neurons in a manner similar to ischemic SD induced by OGD. Can the actions of PLTX provide insight as to how ischemia induces SD? We found that PLTX superfusion of brain slices induces a PLTX depolarization (PD) particularly in the neocortex detected by LT imaging and exhibiting characteristics indistinguishable from OGD-induced SD. Simultaneous field recording and imaging confirmed that the increased LT front was a collective neuronal depolarization within the recorded region. Whole-cell patch recording of pyramidal neurons revealed a rapid, dose-dependent depolarization to zero millivolts similar to SD. Furthermore, as imaged using 2-PLSM, bath PLTX treatment evoked significant pyramidal cell body swelling and dendritic beading with the resultant disappearance of dendritic spines. This neuronal dysmorphia was indistinguishable from OGD-induced injury. LT imaging in brainstem slices revealed that PD affected the same brainstem structures in a similar manner to OGD-induced SD. Finally, the SD-delaying drugs dibucaine (Dib) and carbetapentane (CP) were found to significantly delay PD onset, similar to these two drugs’ delaying effects of OGD-induced SD onset. Thus, we found several striking parallels between the actions of simulated ischemia and of PLTX upon brain slices. Taken together, the findings suggest that an endogenous PLTX-like molecule is released by brain tissue to induce and perpetuate SD during ischemia.

Slice exposure to 10 minutes of 10 nM PLTX reliably evoked SD in the neocortex within 5 to 6 minutes, replicating the well documented findings in our lab’s studies of OGD (Andrew et al. 1999; Douglas et al. 2011; Jarvis et al. 2001a; Obeidat et al. 2000a) and by others (Risher et al., 2012). A focal SD initiation site transformed into a wave front of elevated LT within 10 min, followed by decreased LT in the wake of SD in neocortex, striatum, hippocampus and/or thalamus. Low or no LT changes developed in brainstem gray matter (Brisson et al. 2013, 2014b; Brisson and Andrew 2012). The superficial layers of the tectum where SD could occur were exceptions.

We also measured the negative voltage shift in the extracellular field potential when an OGD-induced propagating wavefront of increased LT passed the recording electrode. Replicating previous studies(Jarvis et al. 2001b; Joshi and Andrew 2001; Obeidat et al. 2000b), this shift coincided with the increased LT front as it passed the recording electrode. It represents a collective depolarization of neurons near the electrode. The consistent recording of a negative shift associated with the LT wavefront demonstrates that it represents SD propagation. This provided us with a standard experimental condition to compare with our novel palytoxin observations.

In addition, 2-PLSM imaging of neocortical pyramidal neurons in layers II/III and V after 10 minutes of OGD reproduced our previous observations of neuron injury in the form of neuron cell body swelling and the loss of dendritic spines with dendritic beading (Davies et al. 2007; Douglas et al. 2011; Jarvis et al. 2001a). Light scattering by dendritic beading accounts for the decreased LT that develops in gray matter regions that had conducted the earlier SD wavefront. This sequence of high, then lower, LT is recapitulated by PLTX treatment evoking PD.

### Palytoxin Evokes Responses Indistinguishable from Spreading Depolarization

When PLTX at concentrations of 1, 10 or 100 nM were bath-applied to coronal brain slices, we observed one or more robust spreading fronts of increased LT in the neocortex, striatum, hippocampus and/or thalamus. The image sequences were identical to SD induced by OGD. The neocortex was the most consistent region to depolarize when exposed to PLTX or to OGD. A higher superfused concentration of PLTX induced depolarization earlier and more consistently. A 10 nM PLTX superfusion induced depolarization in the neocortex at ∼4 minutes while a 1 nM PTX superfusion displayed an onset of ∼7 minutes, indicating a concentration-dependent onset latency for PD. During 10 or 1 nM PLTX superfusion, the neocortex and hippocampus depolarized earlier and more consistently under 10 nM PLTX. Although the striatum and hippocampus displayed an earlier onset in response to 10 nM PLTX, the neocortex and thalamus showed similar vulnerability to 10 nM PLTX and to OGD. Neocortical gray superfused with 1 nM PLTX depolarized between 3-10 minutes while 10 nM PLTX onset was 3-5 minutes, so we opted to use 10 nM for the remainder of the study. Note we found that even concentrations as low as 40 to 50 <u>pM</u> PLTX could evoke SD-like depolarizing fronts (not shown).

In the neocortex, depolarization induced by 10 nM PLTX superfusion displayed a peak ΔLT similar to OGD-induced SD. This PLTX concentration showed a subsequent greater minimum ΔLT with a mean -25% compared to the -13% measured after OGD exposure. This suggests greater acute injury induced by PLTX vs OGD. However Brisson et al. (2014) reported a reduction of -25 to -30% ΔLT in the neocortex after OGD exposure which corresponds with our PLTX value here. Also, dendritic beading may take longer than the mean 23 minutes we spent imaging slices under OGD. Douglas et al. (2011) found -25% ΔLT after a 10 min recovery from 10 minutes OGD in control aCSF. We conclude that the range of acute injury by OGD or PLTX are comparable. Nevertheless, 10 nM PLTX induced consistently and slightly greater cell body swelling of pyramidal neurons (Fig 12).

Previous work by others showed that PLTX depolarizes the membrane potential of skeletal, cardiac and smooth muscle well as spinal cord tissue and nerve fibers (Rossini & Bigiani, 2011). Our intracellular recordings from pyramidal neurons show that bath application of as little as 2 nM PLTX causes marked depolarization; 50 nM evokes complete depolarization in an SD-like fashion. Moreover, the depolarization can propagate as an increased LT front in higher brain regions and initiates earlier at higher PLTX concentrations. This is not unexpected given that PLTX converts the Na^+^/K^+^ pump into a channel open to Na^+^ and K^+^ cations (Artigas and Gadsby 2002, 2004, 2006; Gadsby et al. 2009a; Hilgemann 2003; Hirsh and Wu 1997; Horisberger 2004; Rossini and Bigiani 2011a). The molecular interaction between PLTX and the pump disrupts ion gradients, which in turn perturbs function of other ion-transport systems such as voltage-gated Ca^2+^ channels, Na^+^/Ca^2+^ exchangers and the Na^+^/H^+^ antiport. These downstream events further impair ion equilibria (Rossini and Bigiani 2011a) and, we propose, may also occur during ischemic SD.

### The Negative Shift Confirms a Population Depolarization in Slices

A negative shift of the DC potential accompanying spreading depression in the cerebral cortex was first described by Leao as a surface negative wave or negative “slow voltage variation” that coincided with the spreading depression reaching the region of the electrode placed within the pia mater (Leao 1947; Leão 1951; Marshall 1959) imaged the spreading increase in LT induced by OGD while simultaneously recording intracellularly and found when the increased LT front passed the recording electrode, the recorded cell depolarized rapidly. The coordinated depolarization of local neurons during SD leads to a local negative potential extracellularly, which can be detected by a recording electrode in the vicinity. We confirmed the consistent presence of a sudden negative shift concurrent with the PLTX-induced propagating wave (the PD) at the recording electrode in the neocortex. Those negative shift examples recorded during PD in the neocortex were similar in waveform to the negative shifts we obtained from OGD-induced SD recordings in layer CA1 of the hippocampus. Their amplitudes were indistinguishable. The consistent negative shift as the increased LT wavefront passes the recording electrode indicates a synchronized depolarization of the local neurons induced by PLTX.

### Why are Neurons so Vulnerable to Ischemia orPLTX?

The vague reason commonly cited is that neurons have a higher metabolic rate than other brain cells, but why? We propose that an important reason is the high number and density of NKA complexes on neurons combined with the expansive *surface area* of each neuron. A typical cell has a surface area of 100 to 300 μm^2^ (a human red blood cell is 134 µm^2^) whereas a motoneuron, for example, is 20,000 to 70,000 μm^2^ (P.K. Rose, personal communication). The pump density is 16X cardiac muscle cells and 37X striatal muscle and it has been estimated that there are 50 to 90 pump molecules per dendritic spine (Blom et al., 2016). Imagine then, these high-density transporters opening to elevate Na^+^ and K^+^ conductivity by orders of magnitude.

PLTX has been studied in concert with ouabain and other cardiac glycosides to understand it’s molecular action on the Na^+^/K^+^ pump (Habermann and Chhatwal 1982; Ozaki et al. 1985; Rose and Valdes 1994) as compared to simple pump blockade by ouabain (which also triggers SD). In sharp contrast, PLTX “interferes with the normal strict coupling between inner and outer gates of the pump controlling the ion access to the Na^+^/K^+^-ATPase, allowing the gates to be simultaneously open” (Rossini and Bigiani 2011a). Like OGD, PLTX effects on the NKA are not affected by Na+ channel blockers. In fact, ouabain and other cardiac glycosides act as binding competitors with PLTX. The NKA was identified as an affinity target of PLTX, and that the PLTX binding site on the NKA is coupled to, but not identical with, that of ouabain (Habermann and Chhatwal 1982). Specifically, PLTX binds to the extracellular portion of alpha subunit of NKA on the plasma membrane (Rossini and Bigiani 2011a).

### 2-PLSM Confirms Similar Neuronal Injury Induced by OGD or Palytoxin

Using 2-PLSM, we imaged acute neuronal injury evoked by OGD or PTX on layer II/III pyramidal cells in real time. Like OGD (Andrew et al. 2007), 10 minutes of bath PLTX (1 to 100 nM) caused cell body swelling as well as and the loss of dendritic spines during dendritic beading. This confirms what we detected with LT imaging following PD: the swelling corresponds to the elevated LT front and the decreased LT that follows corresponds to the generation of dysmorphic dendritic beads(Risher et al. 2011b). Our findings of OGD-induced neuronal swelling replicate several previous studies in our laboratory using 2-PLSM (Brisson et al. 2014b; Douglas et al. 2011). This swelling-beading sequence is also observed in *vivo* during intact neocortical ischemia (Brown et al. 2008; Kirov et al. 2020b; Murphy et al. 2008b; Risher et al. 2010).

So, as with +10 minutes of OGD in slices, PLTX leaves pyramidal neurons permanently damaged in the wake of the propagating depolarization it evokes, with dendritic beading and cell body swelling serving as the hallmarks of acute neuronal injury (Jarvis et al. 2001a; Obeidat et al. 2000a; Tanaka et al. 1999). It also drives home the point that ischemia can be as deadly as this potent toxin. It is firmly established that PLTX at even picomolar concentrations can open the NKA to form a channel that allows Na^+^ and K^+^ to flow down their concentration gradients, evoking a strong inward current (Artigas and Gadsby 2002, 2004, 2006; Gadsby et al. 2009a; Hilgemann 2003; Hirsh and Wu 1997; Horisberger 2004; Rossini and Bigiani 2011a). Given the tens of thousands of NKA molecules in a neuron’s plasma membrane, this current would be formidable even if the conductance of each channel is relatively small (Gadsby et al. 2009a). The simultaneous opening of the Na^+^ and K^+^ gates causing the high conductance is both molecularly “subtle” and reversible (Gadsby et al. 2009). Just the right PLTX concentration allows the channel to flicker between open and closed states (Artigas and Gadsby 2002; Gadsby et al. 2009a). A convincing experiment would involve evoking a PLTX-like channel conductance with OGD using micromolar concentrations of ouabain to inhibit channel opening. The opening of PLTX-like channels by OGD (with other Na^+^ and K^+^ channels blocked) would provide direct evidence that SD, generated by ischemia, converts the Na^+^/K^+^ ATPase into an open channel (Gagolewicz et al., 2016). Our data presented in the current study supports this scenario.

### Brainstem ΔLT Imaging Reveals that PLTX Affects the Same Structures as OGD

Previous LT imaging studies of SD in our laboratory found that some brainstem structures are more susceptible to OGD-SD. These include the superficial layers of the superior and inferior colliculi, tegmental nuclei and periaqueductal gray (PAG) area. However, most of the ventral brainstem regions displayed minimal changes in LT following OGD (Brisson et al. 2014b). In the present study, we imaged brainstem regions with stronger illumination to increase resultant ΔLT values in those regions and confirmed a spreading signal in the brainstem dorsal structures noted above. Also, upon PLTX application to brainstem slices, these same structures displayed sensitivity to PLTX. Both PD- and SD-associated increases in LT were followed by decreased LT, often only covering part of the structure imaged. In general, brainstem regions were less damaged by PLTX than higher brain regions, as also observed with OGD (references below). So the selective response of structures within the brainstem to PLTX mimicked the selectivity we observed with OGD. In contrast to higher gray matter, the “lower” hypothalamus and brainstem demonstrate greater resistance to depolarization and better survive global ischemia in patients as follows sudden cardiac arrest (Brisson et al. 2013, 2014b; Brisson and Andrew 2012), possibly accounting for the persistent vegetative state in humans. Patch recordings of neurons in the higher brain display sudden SD onset with a rapid depolarization to near zero millivolts, immediate AP inactivation, and irreversible swelling and dendritic beading of pyramidal neurons in the higher brain (Brisson et al. 2013, 2014b; Brisson and Andrew 2012; Davies et al. 2007). In contrast in the higher brain slices, there is a long latency to SD onset with a gradual depolarization and very slow AP inactivation. Furthermore, the depolarization in lower brain neurons may not reach zero millivolts, and neurons always recover in control aCSF (Brisson et al. 2013, 2014b; Brisson and Andrew 2012). Some studies have shown SD in brainstem regions, but only after severe TBI or in very young rodents(Aiba and Noebels 2015; Richter et al. 2012).

Our finding that the brainstem shows some resilience (relative to higher brain) to such a deadly pump poison like palytoxin demonstrates an additional similarity between the action of this drug and the action of both ouabain and ischemia (Balestrino 1995; Brisson et al. 2014b). Andrew and colleagues have proposed that higher expression of the 1α3 pump isoform by brainstem neurons confers neuroprotection. Our data (Andrew et al. 2017) (Lowry et al. 2020) indicate that higher brain regions strongly express the 1α1 isoform, while lower brain regions express a higher proportion of the 1α3 isoform. Whereas 1α1 is important in maintaining the ionic gradient during basal conditions, 1α3 is vital in maintaining the gradient in non-basal conditions such as a high [Na^+^]_i_ (Azarias et al. 2013). The 1α3 has a greater affinity for ATP relative to 1α1 and should therefore be more resistant to hypoxia and ischemia (Blanco, 2005; Blanco & Mercer, 1998). In addition, studies have shown that 1α3 isoform is required for the rapid restoration of large transient increases in [Na^+^]_i_ and has a lower Na^+^ affinity than 1α1 (Azarias et al. 2013; Munzer et al. 1994; Zahler et al. 1997). So the 1α3 isoform functions more efficiently under ischemic conditions. If the pump channel is more resistant to opening in the 1α3 form, it could explain why PLTX and ischemia are less damaging to brainstem neurons.

### SD Delay by Dibucaine or Carbetapentane

In this study, both 1.0 and 10 µM Dib as well as 30 µM CP were effective in significantly delaying PD and SD. Given the potent action of PLTX, 10 µM Dib was used initially in the current study to assess whether it could delay PD. Once we observed a significant delay, we pretreated neurons with 1 µM Dib, also observing a delay in PD latency. Unlike 10 µM, 1 µM Dib minimally affected normal functioning of neocortical neurons, preserving the evoked field potential amplitude at ∼90% of its initial value (Douglas et al. 2011). Slice pretreatment with 30 µM CP also delayed PD. At 10-50 µM, CP pretreatment for 30-35 minutes delayed OGD-induced SD in neocortical slices (Anderson et al. 2005). At 10 µM, CP (30 minutes) blocked or delayed SD without altering the evoked synaptic response in layers II/III in the neocortex and in layer CA1 of hippocampal slices (Anderson et al. 2005). Our study confirmed these previous studies that found 1-10 µM Dib or 10-50 µM CP pretreatment were effective in delaying SD induced by OGD. Here we also show that these drugs likewise effectively delay PLTX-induced depolarization.

Ouabain has been reported to inhibit PLTX effects by preventing it from binding to NKA because they share similar, but not identical, binding sites (Kim et al. 1995; Wang and Horisberger 1997). CP and dibucaine delay SD by decreasing neuronal excitability but in different ways. Dibucaine raises action potential threshold by blocking the associated sodium channels, while CP reduces excitability by increasing the slow afterhyperpolarization (White et al. 2012), which arises in many neurons when they fire repetitively (Larsson 2013).

### Potential Significance of Our Findings

This study provides evidence that PLTX evokes a strong SD-like event in higher brain gray matter with several properties strikingly similar to SD elicited by ischemia-like conditions. It is well established that PLTX converts the Na^+^/K^+^ pump into an open cationic channel (Artigas and Gadsby 2002, 2004, 2006; Gadsby et al. 2009a; Hilgemann 2003; Hirsh and Wu 1997; Horisberger 2004; Rossini and Bigiani 2011a). We show here that such a conversion will promote spreading depolarization in gray matter because of the double jeopardy of pump failure with the generation of inward current driven by simultaneous Na^+^ influx and K^+^ efflux.

A logical scenario is that ischemia evokes SD in the brain by holding open the gates of the Na^+^/K^+^ pump. This would not only hold true for patients undergoing stroke onset, but for the innumerable patients experiencing global ischemia from sudden heart failure or from massive head trauma. In all these clinical situations *recurring* spreading depolarizations in the higher gray matter are known to worsen outcome (Brisson et al. 2014b; Dreier 2011). Could a recurring opening of the Na^+^/K^+^ pump initiate and drive each event? We have recently shown that PLTX evokes repetitive SD events in the locust CNS (Wang et al. 2024).

Before the open NKA itself was considered a potential channel, several studies showed that the PLTX-induced inward current was unaffected by 1 µM TTX (Karaki et al. 1988; Yoshizumi et al. 1991). These data showed that standard voltage-dependent sodium channels were not involved, as is also the case for OGD-SD.

Neurotoxins commonly target ion channels or ion transporters that drive normal physiological function and thereby shut down the nervous system (Wang 2008). Palytoxin may take advantage of the pump’s vulnerability by holding open the channel in a similar manner to ischemia. Overall, our study documents several striking similarities between the effects of PLTX and of OGD throughout brain gray matter. The possibility that the two treatments share a similar molecular mechanism deserves further study because this concept could supply the link between NKA failure and resultant brain depolarization.

## Acknowledgements

Thanks to Ms. Patti Storey for her technical assistance. This work was supported by grants to RDA from the Heart and Stroke Foundation of Canada, the National Science and Engineering Research Council of Canada and a New Frontiers in Research Fund Exploration Award.

## List of Abbreviations

2-PLSM: 2-Photon Laser Scanning Microscopy
aCSF: Artificial Cerebrospinal Fluid
CBV: Cerebral Blood Volume
CBF: Cerebral Blood Flow
CNS: Central Nervous System
CP: Carbetapentane
Dib: Dibucaine
LT: Light Transmittance
Na_+_/K_+_: ATPase, NKA Sodium-Potassium Activated Adenosine Triphosphatase
OGD: Oxygen Glucose Deprivation
PD: Palytoxin Depolarization
PID: Peri-Infarct Depolarization
PLTX: Palytoxin
PVS: Persistent Vegetative State
ROI: Region of Interest
SD: Spreading Depolarization
TEA: Tetraethylammonium
TTX: Tetrodotoxin

